# CD49a Identifies Polyfunctional Memory CD8 T cell Subsets that Persist in the Lungs after Influenza Infection

**DOI:** 10.1101/2021.07.30.454373

**Authors:** Emma C. Reilly, Mike Sportiello, Kris Lambert Emo, Andrea M. Amitrano, Rakshanda Jha, Ashwin B.R. Kumar, Nathan G. Laniewski, Hongmei Yang, Minsoo Kim, David J. Topham

**Author notes:** authors contributed equally to this manuscript (Emma C. Reilly, Mike Sportiello).

## Abstract

CD8 T cell memory offers critical antiviral protection, even in the absence of neutralizing antibodies. The paradigm is that CD8 T cell memory within the lung tissue consists of a mix of circulating T_EM_ cells and non-circulating T_RM_ cells. However, based on our analysis, the heterogeneity within the tissue is much higher, identifying T_CM_, T_EM_, T_RM_, and a multitude of populations which do not perfectly fit these classifications. Further interrogation of the populations shows that T_RM_ cells that express CD49a, both with and without CD103, have increased and diverse effector potential compared with CD49a negative populations. These populations function as a one-man band, displaying antiviral activity, chemokine production, release of GM-CSF, and the ability to kill specific targets *in* vitro with delayed kinetics compared with effector CD8 T cells. Together, this study establishes that CD49a defines multiple polyfunctional CD8 memory subsets after clearance of influenza infection, which act to eliminate virus in the absence of direct killing, recruit and mature innate immune cells, and destroy infected cells if the virus persists.

**Contribution to the field:** Protection from previously seen infections requires specialized immune memory cells properly positioned throughout the body to combat the newly invading pathogen. In the case of re-exposure to influenza virus, CD8 T cells resident within the respiratory tract (T_RM_) are critical for eliminating the virus. Previously, T_RM_ were viewed as mostly homogenous, with a limited range of immune functions. In this study, lung T_RM_ were compared with circulating memory CD8 T cells transiently present within the lung, to define the breadth of their effector capabilities. Using T_RM_ defining surface proteins CD49a and CD103 to identify different memory CD8 T cell subsets, gene and protein expression were evaluated. In addition to demonstrating higher levels of diversity than previously reported, multiple polyfunctional subsets were identified. This polyfunctionality was primarily associated with cell populations expressing CD49a, and these cells produced multiple antiviral factors, chemokines to recruit other immune cells, a growth factor associated with improved antigen presenting cell function, and cytolytic granules. Functional assays further demonstrated killing of target cells by T_RM_. This study paints a more holistic, complete picture of the phenotype and functions of lung CD8 T cells after viral infection, revealing CD49a as a marker of cells with high effector capacity.

## Introduction

CD8 T cell memory is critical for host protection from previously encountered and related pathogens[1; 2]. After the primary exposure to antigen, multiple classes of memory T cells develop, including circulating (T_CM_ and T_EM_) and resident (T_RM_) populations[3] [4] [5]. T_RM_ are heralded for their ability to rapidly respond upon instances of reinfection, limiting disease severity and improving survival [6] [7] [2; 8]. They are predominantly observed at epithelial barrier surfaces of non-lymphoid tissues, maintained in close proximity to the cells often targeted by viruses ([8] [2]). This memory population has been described in many organs including, but not limited to skin, intestines, lung, female reproductive tract, salivary glands, and tonsils ([9; 10; 11; 12; 13; 14]).

T_RM_ express a range of surface markers, which promote their persistence within the site and facilitate direct interaction with the tissue[15; 16] [17]. These include CD69, an S1PR1 antagonist, CD103/integrin β7, an integrin that attaches to the E-cadherin junction protein between epithelial cells, and CD49a/CD29, a collagen binding integrin [18; 19] [20; 21]. The requirement for these surface receptors differs between organs, with CD69 playing critical roles in the development and maintenance of kidney T_RM_, but with less reliance in other sites including lung [15; 22]. CD103 contributes to the accumulation of T_RM_ early during the resolution phase, with minimal effects on long term persistence in the intestines, salivary glands, and lung, though maintained expression may be critical in other organs or for functions other than retention within the tissue [12; 15; 23; 24]. A dependence on CD49a for the survival of T_RM_ and subsequent protection has been demonstrated in the pulmonary system and intestines, and a role for increased effector capacity has been shown in both skin and tumor T_RM_[13; 25; 26; 27]. The observed heterogeneity in the surface phenotype of T_RM_ in different organs is likely driven by microenvironmental differences and the composition of the immune response elicited by the infecting pathogen.

One of the main immune contributors to these differences is the cytokine milieu that results from infection and resolution. TGFβ is known to mediate expression of CD103 and CD49a and a role for IL-12 in promoting surface CD49a while limiting CD103 has been demonstrated *in vitro*[28; 29]. While it makes sense that differences exist between organs, variations in features including the immune cell composition, cytokine levels and antigen load also differ within discrete niches in a given organ. With this understanding, we hypothesized that the population of cells in the lungs referred to as T_RM_, which arises after resolution of influenza A virus infection, actually represents a heterogenous mix of multiple subpopulations. To address this, surface phenotyping and RNAseq analyses were performed after resolution of disease in the presence or absence of stimulation. Within these memory populations, the question still remained as to whether certain populations display higher effector capabilities. Using a combination of RNAseq and flow cytometry, this study set out to determine which memory cell(s) offer the most potent protection potential during secondary encounter with antigen.

## Materials and Methods

### Mice

All mice were housed in university-approved microisolator cages, within a pathogen-free facility. C57BL/6J mice (Jackson Laboratories) used for experiments were infected at 8-10 weeks of age. This study was carried out in strict accordance with the recommendations in the *Guide for the Care and Use of Laboratory Animals* as defined by the NIH[30]. Animal protocols were reviewed and approved by the Institutional Animal Care and Use Committee of the University of Rochester. All animals were housed in a centralized and Association for Assessment and Accreditation of Laboratory Animal Care accredited research animal facility that is fully staffed with trained husbandry, technical, and veterinary personnel.

### Virus and Infection

Mice were anesthetized with 3,3,3-tribromoethanol (Avertin). Upon verification of sedation, mice were placed in the supine position and infected intranasally with 10^5^ EID_50_ of HKx31 human influenza A virus or 3×10^3^ EID_50_ HKx31-OVAI expressing the OVA^257-264^ SIINFEKL peptide in the stalk of the neuraminidase in 30uL volume. Mice were observed until they recovered from anesthesia and monitored daily for weight and overall morbidity.

### Tissue harvesting and processing

Mice were anesthetized with Avertin and upon sedation mice were injected intravenously with 0.2ug labeled CD45 antibody in 100uL sterile 1x PBS[31]. For figures 1-8, after 3 minutes, the peritoneal cavity was opened, the aorta was cut, and the lungs were harvested. For figure 9, bronchoalveolar lavage (BAL) and cardiac puncture were performed prior to opening the peritoneal cavity and the organs were harvested in the following order: spleen, MLN, lung. BAL and blood cells were spun down at 300xg for 6 minutes, lysed with 0.5mL or 3mL, respectively 1x ammonium-chloride-potassium (ACK) lysis buffer for 5 minutes at room temperature, and washed with 14mLs 1x PBS with 1% FBS (PBS serum). Spleen and MLN were dissociated using the frosted ends of frosted glass slides. These samples were put through 100um filters and spun at 300xg for 6 minutes, followed by ACK lysis with 3mL or 0.5mL, respectively. Samples were washed with PBS serum. Lungs were separated into the left and right lobes prior to dissociation in a Miltenyi Biotec C tube containing 2mL [2mg/mL] collagenase II (Worthington) in RPMI with 8% FBS (RPMI serum) using the Miltenyi Biotec GentleMACs Lung01 program. Collagenase II volume was increased to a total of 5mLs and samples were incubated upside down with gentle shaking at 37C for 30 minutes. Lung samples were further dissociated using the GentleMACs Heart01 program. These samples were filtered through 100um strainers and run on a discontinuous 75/40% Percoll (Cytiva) gradient. Cells were harvested from the interface and washed with PBS serum for immediate staining or RPMI serum for in vitro stimulation.

**Figure 1.**
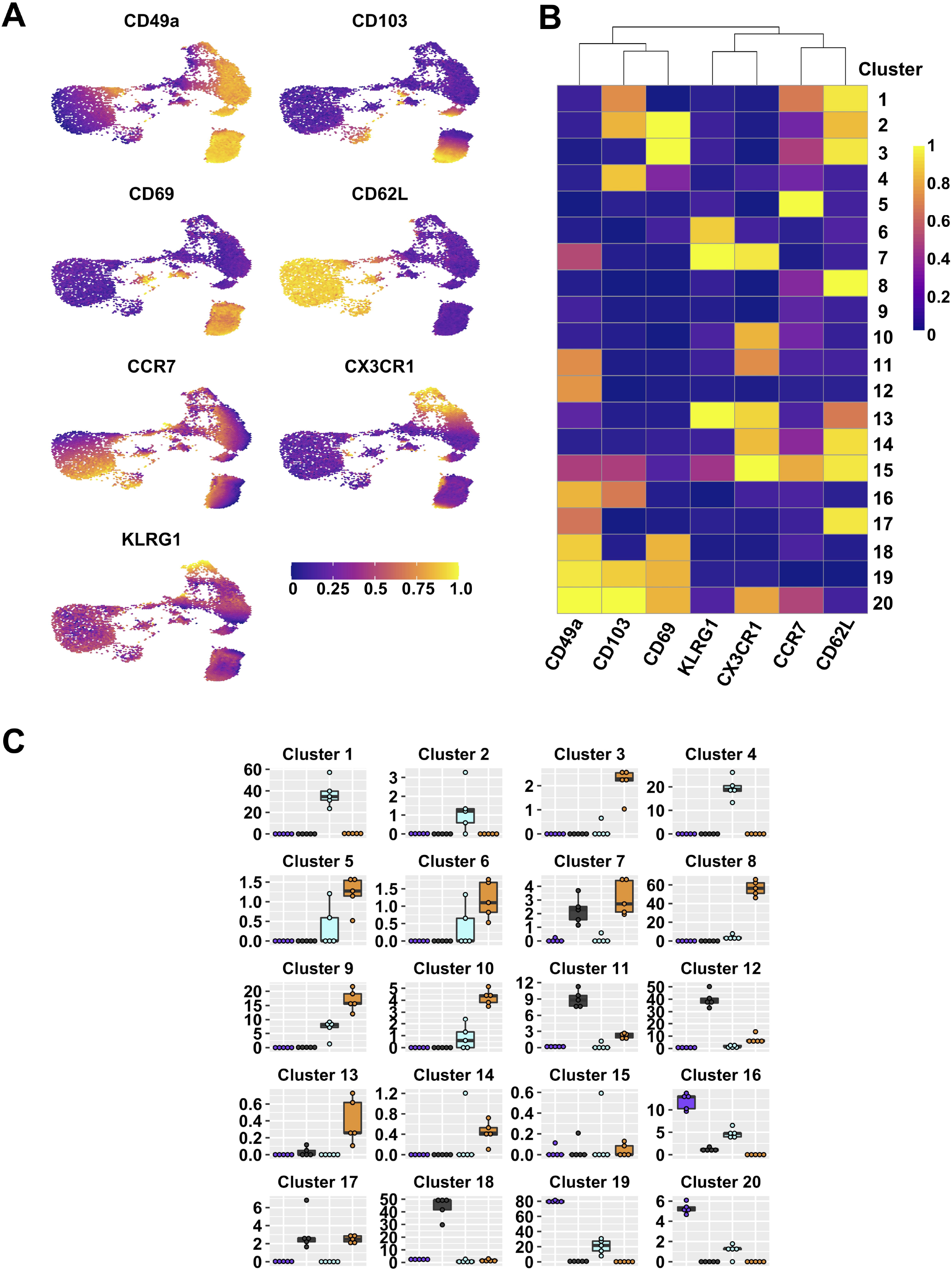
Integrin subsets are heterogenous based on classical memory T cell markers. UMAPs displaying expression level of different effector and memory markers on lung tissue CD44^+^ CD8 T cells (A). FlowSOM clusters based on expression of indicated markers; heat map indicates range scaled (0-1) median expression levels (B). Frequencies within lung tissue DP, CD49a, CD103, and DN populations that contribute to each cluster (C). * p<0.05 based on a repeated measures one-way ANOVA, followed by *post-hoc* testing comparing groups to DN using Dunnett’s multiple comparisons test. Data is from 1 representative experiment of 2 independent experiments with n≥ 3 mice/experiment.

### *In vitro* stimulation

For in vitro culture, 96-well dishes were used. Wells were pre-coated with αCD3 and αCD28 purified antibodies at [5ug/well] in 1x PBS where indicated. NP and PA peptides were added at 1ug/mL. GolgiPlug™ (Brefeldin A) and GolgiStop™ (Monensin) were used as indicated by BD Biosciences (1uL/mL and 4µL/6mL respectively) for 6 hours prior to staining. Anti-Lamp-1 antibody was added as indicated by manufacturer (4µL/well for 4 hours) prior to staining. GM1006 MMP inhibitor was used as indicated by the Millipore Sigma (25µM) for FasL staining.

### *In vitro* killing

1×10^6^ GFP OT-I splenocytes were transferred IV to naïve C57BL/6 males aged 8-10 weeks. Mice were infected with HKx31-OVAI the following day. Cells were harvested from the lung tissue of IV labeled mice at day 21 or day 28 and processed to a single cell suspension. Cells negative for IV labeling were sorted based on integrin phenotype into CD49a single positive, CD49a CD103 double positive, and double negative groups. EL4 cells were counted and incubated at 1×10^6^ cells/mL in RPMI with CellTrace™ Violet as recommended by the company, and 10ug SIINFEKL peptide or equivalent amount of vehicle (dH_2_O) for 45 minutes at 37C [32]. All cells were washed three times with volumes of 15 mL RPMI serum. Sorted cells and EL4 cells were plated at a 1:1 ratio in U-bottom plates for 11hours. An additional 5ug of SIINFEKL peptide or an equivalent amount of vehicle were added to wells for 2 hours prior to harvest. Cells were transferred to V-bottom dishes, washed with cold PBS, and resuspended in 100uL Annexin V Binding Buffer containing 5uL Annexin V per sample, and transferred to 5mL FACs tubes. After 30 minutes, 200uL cold Binding Buffer added to each tube, and samples were evaluated by flow cytometry, in pairs of SIINFEKL and vehicle controls per sample.

### Flow cytometry

Cells were spun down at 800xg for 5 minutes in 96-well V-bottom plates. Supernatant was flicked off and cells were resuspended in a master mix of FC-block, surface antibodies, and Aqua Fixable Viability Dye. Antibodies were added at 1:200 unless otherwise indicated and viability dye was added as suggested by manufacturer. Cells were incubated in the dark at room temperature for 30 minutes. The BD intracellular staining kit was used as indicated including recommended washes. Briefly, cells were fixed and permeabilized in 1x Fix/Perm for 25 minutes, followed by washing and intracellular staining in Permwash at room temperature for 30 minutes. After washes in Permwash and PBS serum, cells were resuspended in PBS serum and data was captured on an 18 color LSRFortessa with 5 laser lines (Blue, Green, Red, Violet, UV). All antibodies were purchased from BD Biosciences, Biolegend, or Invitrogen (S Table 1).

### Bulk RNA sequencing

For each experiment, lungs were harvested from three pools of 5 mice each and processed as described. Cells were negatively enriched for CD8 cells using the Miltenyi Biotec mouse CD8^+^ T Cell Isolation Kit (Miltenyi Biotec, Bergisch Gladbach, North Rhine-Westphalia). For the restimulated group, cells were plated on αCD3/28 coated plates for 5 hours. CD8 T cells were sorted for live, singlet, IV^neg^CD45IV^neg^CD8^+^CD44^+^ cells followed by the four CD49a CD103 quadrants into RPMI + serum + penicillin/streptomycin. Cells were spun down and resuspended in RNAeasy buffer containing beta mercaptoethanol. Total RNA was isolated using the RNeasy Plus Micro Kit (Qiagen, Valencia, CA). RNA concentration was determined with the NanopDrop 1000 spectrophotometer (NanoDrop, Wilmington, DE) and RNA quality assessed with the Agilent Bioanalyzer 2100 (Agilent, Santa Clara, CA). 1ng of total RNA was pre-amplified with the SMARTer Ultra Low Input kit v4 (Clontech, Mountain View, CA) per manufacturer’s recommendations. The quantity and quality of the subsequent cDNA was determined using the Qubit Flourometer (Life Technologies, Carlsbad, CA) and the Agilent Bioanalyzer 2100 (Agilent, Santa Clara, CA). 150pg of cDNA was used to generate Illumina compatible sequencing libraries with the NexteraXT library preparation kit (Illumina, San Diego, CA) per manufacturer’s protocols. The amplified libraries were hybridized to the Illumina flow cell and sequenced using the NextSeq 550 sequencer (Illumina, San Diego, CA). Single end reads of 75nt were generated for each sample. Reads were aligned with STAR-2.7.0 and reads quantified with Subread-1.6.4. DESeq2-1.26.0 was used to generate normalized count matrices and to assess differential expression. IHW-1.14.0 was used to perform independent hypothesis weighting to correct p values. Final, adjusted, weighted p values were used to assess differential expression. A cutoff of p<0.05 was used. Enrichr-2.1 was used for gene set enrichment analysis to assess for enrichment of pathways from the Kyoto Encyclopedia of Genes and Genomes (KEGG), for which a p<.05 was used. EnhancedVolcano-2.1 was used to plot volcano plots. ggplot2-3.3.2 was used for a variety of plots. Pheatmap-1.0.12 and gplot-3.1.1 were used for heatmap plotting. A full list of packages, version numbers, and software citations can be found on github: https://github.com/tophamlab20.

### Statistical analysis of flow cytometry data

Statistics were performed using Prism analysis software (GraphPad 9). Percentage data was log2 transformed prior to statistical analysis. Repeated measures one-way ANOVA were performed with Greenhouse-Geisser correction, followed by *post-hoc* testing comparing groups to DN using Dunnett’s multiple comparisons test. When comparing *in vitro* treatment to control, a paired T-test was performed on log2 transformed data. MFI data were normalized to the DN population per mouse. Only samples with cells in all four quadrants were included. Significance was defined as a p-value<0.05 (or adjusted p-value when appropriate).

### Results

After influenza infection, the lung is home to multiple types of CD8 T cells. Based on the established paradigm, CD8 T cells in the tissue (IV-label negative) constitute circulating T_EM_(CD44^pos^CD62L^neg^KLRG1^neg^CCR7^neg^CX3CR1^pos/neg^) and predominantly non-circulating T_RM_ (CD44^pos^CD69^pos^CD49a^pos^CD103^pos^) [5; 33; 34] [8] [2]. To verify the presence of these two main subsets and determine the level of heterogeneity within each memory type, cells were examined directly *ex vivo* after clearance of influenza virus at day 21 post-infection by flow cytometry. The cells were stained for classical effector and memory surface markers, allowing for the discrimination of T_EFF_ (CD44^pos^CD62L^neg^KLRG1^pos^), T_CM_ (CD44^pos^CD62L^pos^), T_EM_ (CD44^pos^CD62L^neg^KLRG1^neg^CCR7^neg^CX3CR1^pos/neg^), and T_RM_ (CD44^pos^CD69^pos^CD49a^pos^CD103^pos^) [5; 33; 34] [8] [2]. Dimensional reduction and visualization of data from TCRβ^pos^CD8^pos^CD44^pos^ cells was achieved through Uniform Manifold Approximation and Projection (UMAP) (Figure 1A) [35]. Data were subsequently clustered using FlowSOM, resulting in 20 distinct clusters (Figure 1B) [36]. Unexpectedly, some of the cells expressed CD62L, resulting in multiple CD62L^pos^ clusters. While this would be anticipated in the vasculature associated population and lymphoid organs, it is not well established that these cells would also be present in peripheral non-lymphoid tissues, or perhaps suggests the tissue includes some lymphatic structures (Figure 1B and S Figures 1 and 2). Clusters 1 and 2 co-express CD103, along with CCR7 and CD69, respectively. Cluster 3 co-expresses CD69 alone, cluster 13 also stains for KLRG1 and CX3CR1, suggesting that they may represent a transition state between effector and memory. Classical T_EM_ (CD44^pos^CD62L^neg^KLRG1^neg^CCR7^neg^CX3CR1^pos/neg^CD49a^neg^CD103^neg^) are found in clusters 9 and 10, in the absence and presence of CX3CR1 staining, respectively. Clusters 11 and 20 both have CX3CR1, but co-express CD49a alone or CD49a, CD103, and CD69, supporting that some subsets of “T_RM_” phenotype cells may have the capacity to leave the tissue[37]. Alternatively, CX3CR1 can bind its chemokine ligand fractalkine (CX3CL1) expressed on the surface of epithelial and endothelial cells, potentially aiding in micropositioning within different niches throughout the tissue[38]. Finally, cluster 19, expressing the combination of CD49a, CD103, and CD69 contains the cells considered to be classical T_RM_. No subsets expressing only CD103 and CD69 were found, however cluster 4 represents CD103 single positive cells. On the contrary, cluster 18 expresses both CD49a and CD69. Clusters 12 (CD49a only) and 16 (CD49a and CD103) may also indicate CD69 negative T_RM_ subsets.

Since CD69 did not appear to be the defining feature of lung T_RM_, we focused on further subsetting the memory CD8 T cells based on CD49a and CD103 expression. CD49a^pos^CD103^neg^(CD49a), CD49a^pos^CD103^pos^ (<u>D</u>ouble <u>P</u>ositive DP), CD103^pos^CD49a^neg^ (CD103), and CD49a^neg^CD103^neg^ (<u>D</u>ouble <u>N</u>egative DN) were examined for their contribution to each of the twenty clusters (Figure 1C and S Table 2). CD103 was represented across clusters 1 and 4, and based on phenotype, has the potential to function as T_CM_ and possibly CD103 single positive T_RM_. However, similar analysis at three months post-infection demonstrates that the CD103 single positive T_RM_ population does not persist long-term. The DN population is accounted for by a number of clusters (3, 5, 6, 8, 9, and 10), and in addition to the hypothesized T_EM_ phenotype, these cells also represent T_CM_ and effector-like populations. As a whole, these subsets likely represent “circulating” CD8 T cell memory. The CD49a population is predominantly found in clusters 11, 12, and 18, with the potential ability to recirculate based on CX3CR1 expression in cluster 11. DP cells are distributed across clusters 16, 19, and 20, with a small subset expressing CX3CR1 (cluster 20).

The phenotyping data suggest that as a whole, the DN and CD103 subsets represent circulating memory cells and the CD49a and DP subsets are likely the predominantly resident populations. Each memory population is presumed to be capable of performing immune surveillance, and the cells expressing CD49a are highly motile, yet their functions in the homeostatic state are unclear [39]. To further characterize these subsets, non-naive CD8 T cells from the lung tissue (CD45IV^neg^/TCRβ^pos^/CD8^pos^/CD44^pos^) of day 21 infected mice were sorted based on their CD49a/CD103 profile, and bulk RNA sequencing was performed on these four subsets to determine if there were intrinsic baseline differences under homeostatic conditions (S Figure 3). Principle component analysis demonstrated that transcript expression stratifies the CD49a expressing subsets (CD49a and DP) from the CD49a negative populations (CD103 and DN), but it is insufficient to further distinguish the subsets into CD103 positive and negative (Figure 2A). PC1, which accounts for over 40% of the variance is driven in part by *Itga1* (CD49a) which serves as an internal control for sorting (S Figure 4A,B). PCA excluding *Itga1* did not alter the patterns observed suggesting additional genes define these subsets (S Figure 4C-E). Additionally, PC1 shows upregulation of *Ifitm1*, adhesion molecule *Tjp1* (ZO-1), and *Csf1* (S Figure 4A). Since the DP population demarcates the canonical T_RM_ cells and DN are circulating memory, the initial analysis compared these two populations to investigate any transcriptional differences. Using independent hypothesis weighting and adjusted p-values, over 700 genes were found to be differentially expressed when comparing DP to DN populations[40]. Examination of the top 500 genes that displayed differential regulation further supported that the CD103 and DN cells are almost transcriptionally indistinguishable at baseline (Figure 2A-D). CD49a cells also display significantly different transcript levels from CD103 and DN cells, and to a smaller degree compared with DP (Figure 2A-C). In fact, a small subset of genes is uniquely upregulated in CD49a cells than DP cells (Figure 2A-C).

**Figure 2.**
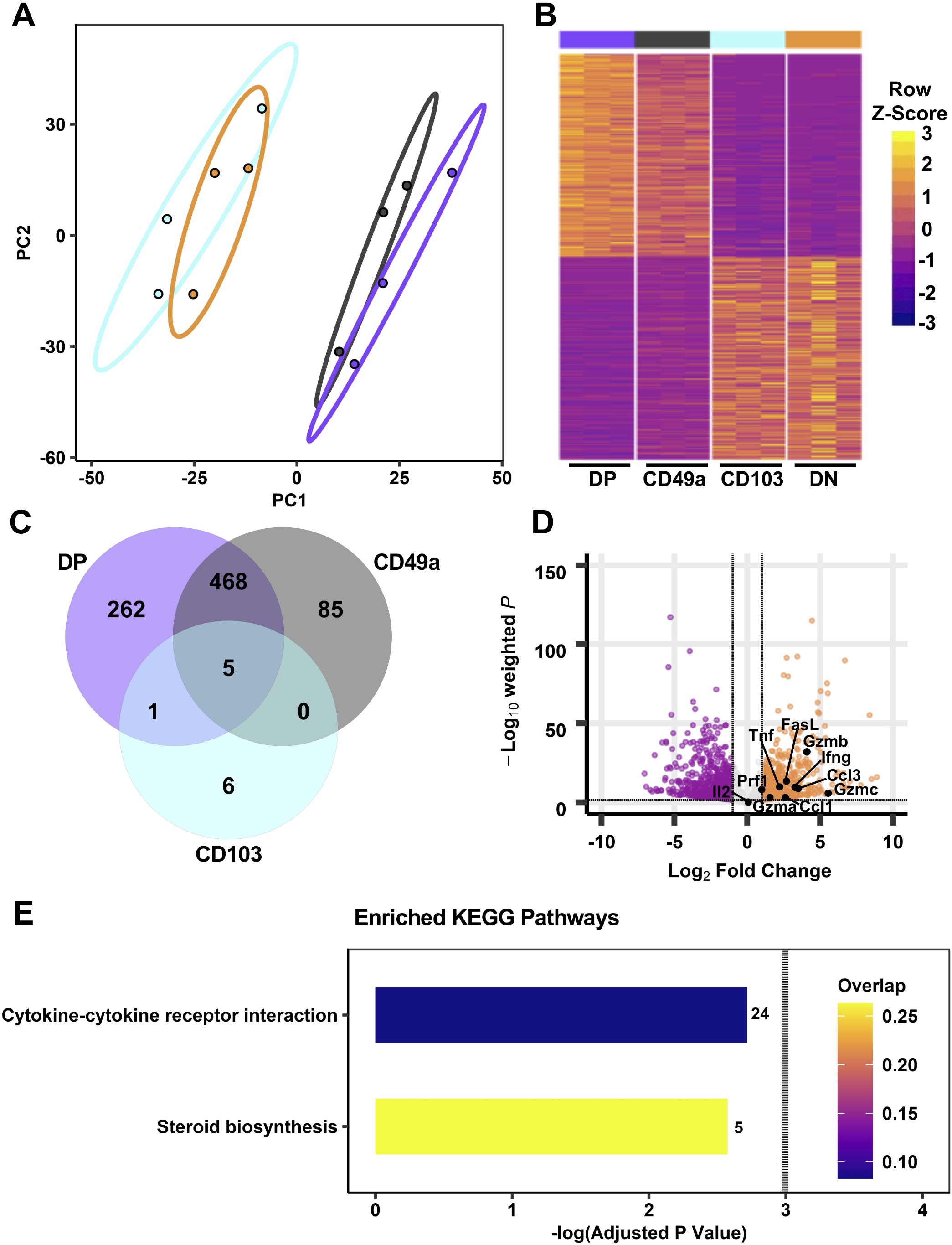
Transcriptional profiles of CD49a expressing memory CD8 T cell subsets are distinct from CD49a negative counterparts at baseline. PCA of transcripts from DP, CD49a, CD103, and DN populations (A) and heat map of the top 500 differentially expressed genes (B). Venn diagram showing the number of genes differentially expressed compared with DN cells (C). Effector genes overlaid on the volcano plot comparing DP to DN (D). Kegg enriched pathways based on a significance of adj p<0.1 with the bar color indicative of the overlap between upregulated genes within the pathway and the number of overlapping genes indicated. Dotted line indicates p=0.05 (E). Data is from three groups of five mice pooled and sorted for integrin phenotype.

To further appreciate the cellular programs which are the foundation for these observed transcriptional differences, Kyoto Encyclopedia of Genes and Genomes (Kegg) pathway gene set enrichment analysis (GSEA) was performed to reveal differentially regulated biological pathways [41; 42; 43; 44]. Despite over 700 genes with differential expression between DP and DN cells, no pathways were significantly different (padj<0.05). Using a less strict level of significance (padj<0.1) two pathways were identified: *cytokine-cytokine receptor interaction*s and *steroid biosynthesis* (Figure 2E). With the goal of defining the population(s) that are most effective at limiting re-infection, the *cytokine-cytokine receptor interactions* pathway was more closely scrutinized. Directed examination identified a number of transcripts that were differentially expressed at baseline. These include: *Tnf, Ifng, Gzma, Gzmb, Gzmc, Prf1*, and the chemokine *Ccl3* (Table 1, S Table 3, and Figure 2D). Protein levels for these effector molecules were quantified by flow cytometry. However, despite an increase in transcript levels, only low levels of granzyme A, perforin, and Lamp-1 were detected (S Figure 5). This lack of correlation between cytokine transcript and protein levels has been seen previously in T_RM_ cells [45; 46]. To further compare with T_RM_ subsets, pathways were compared between the DP and the CD49a populations. However, at baseline, no differentially expressed pathways were identified (Figure 2E and S Table 4).

**Table 1.** Effector genes upregulated between DP and DN memory CD8 T cells at homeostasis. Effector genes that are significantly upregulated in DP cells compared with DN cells, ordered by log2 fold change.

| Gene name | Protein name | Log2 Fold Change | Base Mean | Adj. p-value |
| --- | --- | --- | --- | --- |
| *Gzmc* | Granzyme C | 5.55 | 17 | 1.65E-06 |
| *Gzmb* | Granzyme B | 4.09 | 9,250 | 1.29E-32 |
| *Ccl3* | CCL3 | 3.50 | 831 | 1.99E-09 |
| *Ifng* | Interferon $\gamma$ | 3.28 | 3,595 | 2.86E-10 |
| *Ccl1* | CCL1 | 2.62 | 32 | 8.20E-4 |
| *Tnf* | TNF | 2.23 | 888 | 1.82E-10 |
| *Gzma* | Granzyme A | 1.56 | 2063 | 7.87E-04 |
| *Prf1* | Perforin | 1.00 | 6,979 | 9.12E-09 |

Under homeostasis, the prototypical T_RM_ (DP) and the CD49a population both expressed higher baseline transcript levels for effector genes compared with non-T_RM_ memory. However, it was unclear whether this observation persisted in the different subsets after restimulation. To address this, cells were activated *in vitro* with αCD3 and αCD28, as a mimic for TCR-based stimulation. Briefly, memory CD8 T cells were sorted on day 21 post-infection after 5 hours of *in vitro* stimulation using a similar approach to the unstimulated cells (CD45IV^neg^/CD8^pos^/CD44^pos^ and sorted based on CD49a/CD103 expression). PCA of the bulk RNA sequencing results demonstrates that after stimulation, the transcriptional profiles for all subsets are distinct (Figure 3A, S Table 5, and S Figure 6). Top loadings for the principal component separating CD49a expressing subsets (PC1) from CD49a negative subsets include effector molecule *Ccl1* and junctional protein *Tjp1* (S Figure 6A). Component 1 contributes to over 60% of the variance. The gene encoding CD49a (*Itga1)* again serves as an internal control for the sorting technique used. Nevertheless, the PCA results are nearly identical when CD49a and CD103 are excluded (S Figure 6C-E). The top loadings of PC2, which separates CD103 expressing subsets from CD103 negative subsets also include both effector-associated genes (*Ccl9* and C*x3cr1*) and structural binding proteins including *Cdh4* and *Itgae* (which encodes CD103 and serves as an internal control). The number of differentially expressed genes, when comparing DP to DN increased to over 1,300, with almost 800 genes uniquely upregulated only in the DP cells (Figure 3B-D). GSEA of the differentially regulated genes revealed over 70 upregulated KEGG pathways when comparing DP to DN. The preponderance of pathways uncovered related to effector programs and the top differentially expressed pathway was *cytokine-cytokine receptor interactions* (Figure 3E). Other enriched pathways include *C-type lectin receptor signaling pathway, cell adhesion molecules, natural killer cell mediated cytotoxicity, chemokine signaling pathway*, and *TNF signaling pathway* (Figure 3E and S Table 6). Within the *cytokine-cytokine receptor interaction pathway*, DP showed increased levels of 54/292 genes and specific evaluation of effector associated genes that were upregulated in the DP compared with the DN population identified 17 genes to further pursue (Figure 2D and Table 2). Of note, one of the most upregulated genes in the data set was *Ccl1*, the chemokine associated with recruitment of innate myeloid derived cells through binding its cognate receptor CCR8 [47; 48]. DP cells also displayed increased transcript levels for genes associated with classical antiviral cytokines (*Ifng, Tnf*), T cell survival and activation (*Il2*), chemokines (*Ccl3, Ccl4, Cxcl9*), cell differentiation (*Gmcsf*) and cytolytic components (*Prf1, Gzma, Gzmb,Gzmc, Fasl, Tnfsf10*) (Table 2).

**Figure 3.**
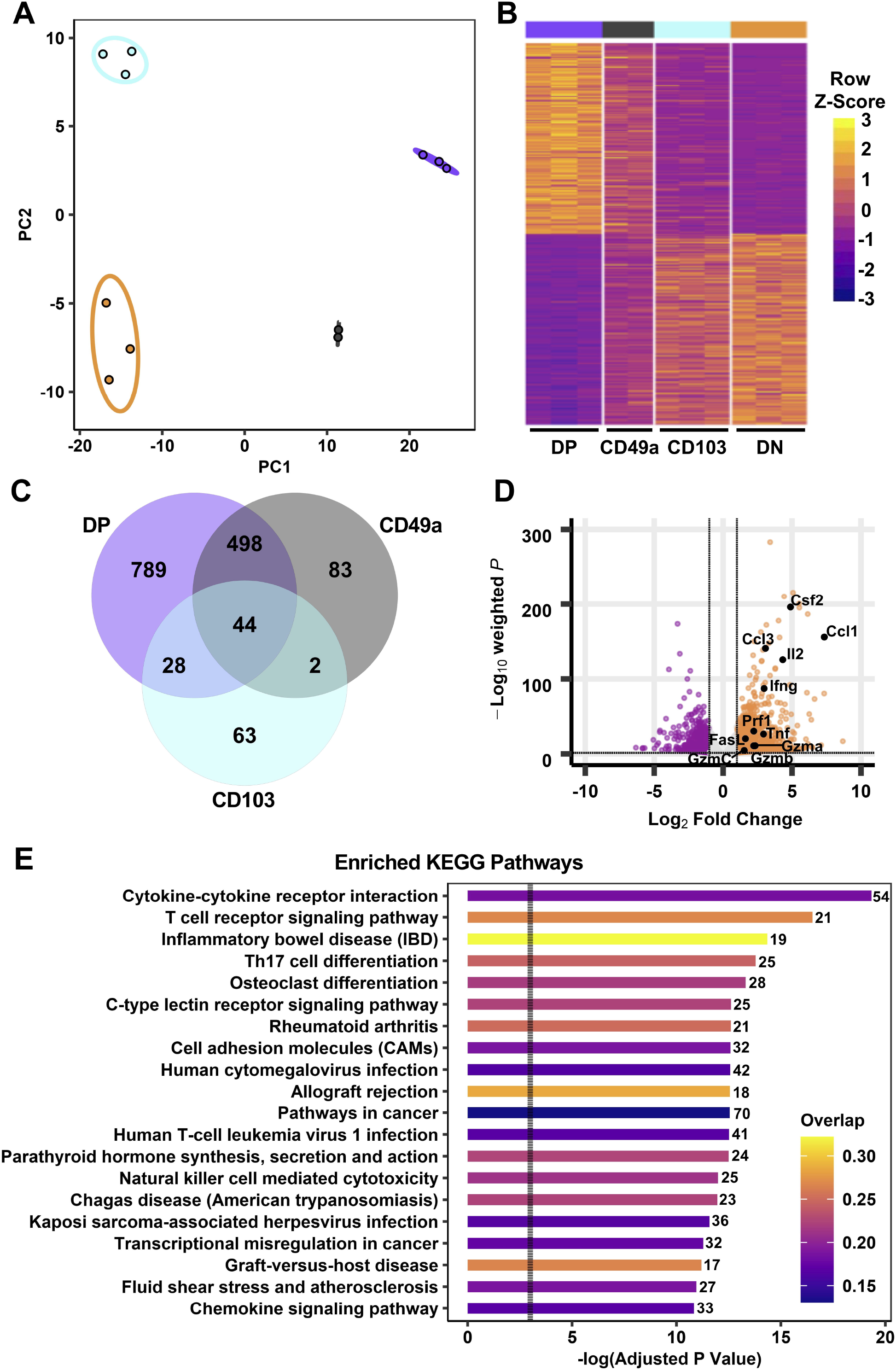
Transcriptional profiles of integrin memory subsets are all distinct after *in vitro* re-stimulation. PCA of transcripts from DP, CD49a, CD103, and DN populations (A) and heat map of the top 500 differentially expressed genes (B) after five hours *in vitro* αCD3/CD28 stimulation. Venn diagram showing the number of genes differentially expressed compared with the DN cells (C). Effector genes overlaid on the volcano plot comparing DP to DN (D). Kegg enriched pathways based on a significance of adj p<0.05 with the bar color indicative of the overlap between upregulated genes within the pathway and the number of overlapping genes indicated. Dotted line indicates p=0.05 (E). Data are from three groups of five mice pooled and sorted for integrin phenotype. One CD49a sample did not reach RNA quality cutoff and was excluded from the analysis.

**Table 2.** Effector genes upregulated between DP and DN memory CD8 T cells after *in vitro* stimulation. Effector genes that are significantly upregulated in DP cells compared with DN cells, ordered by log2 fold change.

| Gene name | Protein name | Log2 Fold Change | Base Mean | Adj. p-value |
| --- | --- | --- | --- | --- |
| *Ccl1* | CCL1 | 7.34 | 37,106 | 1.26E-156 |
| *Csf1* | CSF | 6.14 | 2,939 | 2.48E-187 |
| *Csf2* | GM-CSF | 4.89 | 2,769 | 1.02E-196 |
| *Il2* | IL-2 | 4.33 | 5,369 | 2.35E-126 |
| *Il10* | IL-10 | 4.22 | 460 | 9.37E-51 |
| *Ccl3* | CCL3 | 3.07 | 129,614 | 1.59E-141 |
| *Ifng* | Interferon $\gamma$ | 2.98 | 104,386 | 5.95E-88 |
| *Tnf* | TNF $\alpha$ | 2.93 | 12,511 | 4.02E-27 |
| *Tnfsf10* | TRAIL | 2.76 | 565 | 1.47E-41 |
| *Gzma* | Granzyme A | 2.33 | 2,379 | 1.32E-11 |
| *Prf1* | Perforin | 2.23 | 7,903 | 4.38E-31 |
| *Gzmb* | Granzyme B | 2.22 | 86,942 | 1.66E-11 |
| *Il21* | IL-21 | 1.96 | 144 | 4.39E-07 |
| *Fasl* | Fas Ligand | 1.62 | 2,716 | 5.92E-21 |
| *Gzmc* | Granzyme C | 1.53 | 1,205 | 2.40E-05 |
| *Cxcl9* | CXCL9 | 1.26 | 542 | 7.04E-04 |
| *Il17a* | IL-17A | 1.31 | 100 | 8.06E-04 |

A requirement for T_RM_ derived IFNγ is known to be critical for protection against influenza, so these findings support the contribution of the DP population to the antiviral response [49]. Notably, the CD49a population showed similar increases in effector transcript levels compared with DN T cells, suggesting that they may represent a previously unappreciated critical antiviral T_RM_ subset. In fact, a direct comparison between the DP and CD49a showed that genes were more highly enriched for the *natural killer cell mediated* cytotoxicity pathway in CD49a, suggesting that the cytotoxic components may be further enhanced in the CD49a single positive cells compared with DP. Conversely, the majority of effector genes were not differentially expressed between CD103 and DN, and in some cases displayed higher transcript levels in DN, such as *Ifng*.

To determine whether transcript levels equated to protein expression after re-stimulation, flow cytometry was utilized (Figure 4). Additionally, one focus was evaluating whether T_RM_ are polyfunctional, or if separate sub-populations contribute to only one or two aspects of the antiviral response. To achieve this, T cells were stimulated *in vitro* with αCD3/CD28 in the presence of Golgi and ER transport inhibitors, respectively for 6 hours, and examined by multi-color intracellular flow cytometry [50]. CD8 T cells were surveyed for antiviral cytokines, IL-2, CCL1, and cytolytic components based on Table 2 (Figure 4A). The multidimensional data from the four integrin subsets were concatenated and reduced down to two dimensions using UMAP to generate a similarity map based on the functional molecules examined (TNF, IFNγ, IL-2, Granzyme A, Perforin, Lamp-1, and CCL1) (Figure 4C) [35]. FlowSOM clustering was applied to further interrogate the T cell effector profiles [36]. This approach yielded 18 distinct clusters (Figure 4B). Given the prior understanding of beneficial effector capabilities, focus was put on examining clusters 10, 13, 14, and 16-18, all of which identify polyfunctional subsets. Cells in these clusters are CCL1^+^ with co-expression of IFNγ and TNF, but further separated based on the level of these molecules, as well as staining for perforin, IL-2 and Lamp-1 (Figure 4B,D). Strikingly, when examining the frequencies of these clusters within the integrin subtypes, DP cells had the highest proportions for all of these clusters (Figure 4D and S Table 7). Although less prominent than the DP cells, CD49a cells also displayed marked contributions to these clusters (Figure 4D and S Table 7). In contrast, the vast majority of CD103 and DN cells could be accounted for with clusters 3 and 5, which displayed low levels of perforin and TNF or low quantities of only perforin, respectively. These data support that CD49a and DP identify multiple polyfunctional T_RM_ subsets.

**Figure 4.**
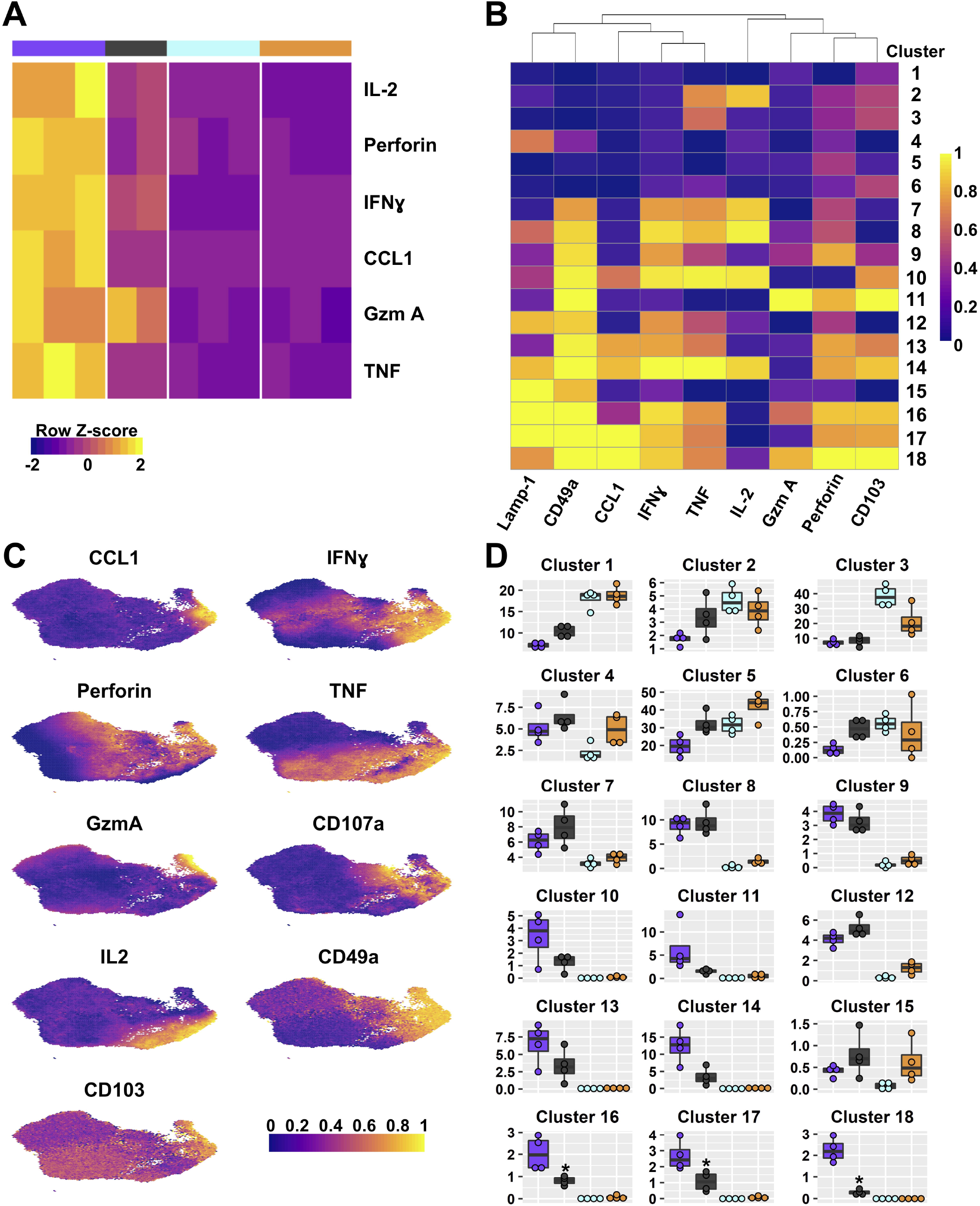
DP and CD49a CD8 memory T cell subsets are polyfunctional. Heat maps show the transcriptional differences between the four T cell subsets (A) and the clusters based on protein staining for Lamp-1, CCL1, IFNγ, TNF, IL-2, Granzyme A, and Perforin after 6 hours of *in vitro* stimulation; heat map (B). UMAPs are based on the effector molecules evaluated (C). Percentage of the integrin subset that is found within each cluster (D). * is p<0.05 based on multiple comparisons ANOVA followed by the Dunnett’s post hoc test comparing to DN. Data shown is one experiment representative of at least three for B-D with an n≥ 3 mice/experiment. DP CD49a CD103 DN

In an effort to further interrogate the presence of effector molecules in the different memory subsets, the frequency, normalized count, and MFI of the positive fractions were quantified for an expanded panel of cytokines, chemokines, a growth factor, and cytotoxic components. Both DP and CD49a cells showed increases in frequencies for IFNγ and IL-2 compared with DN cells, with DP cells also showing increases in the percentage of TNF positive cells (Figure 5A,B). DP and CD49a also had increased MFIs for both IFNγ and TNF compared with both CD103 and DN (Figure 5D). This approach demonstrates not only higher proportions of positive cells, but also supports increased protein levels on a per cell basis in DP and CD49a compared with non-T_RM_ cells (Figure 5B,D). Among the IFNγ^+^ cells, DP and CD49a were the main contributors, comprising approximately 80% of the response (Figure 5C). Despite a strong bias toward CD49a expressing subsets producing more IFNγ and IL-2, an increased percentage of CD103 cells were found to make TNF (as shown in cluster 3) at comparable frequencies with DN, but this was not accompanied by a higher contribution to the TNF^+^ population or a greater amount produced on a per cell basis (Figure 5B,D). To ensure that the outcomes observed with bulk stimulation of CD8 T cells weren’t skewed from that of known influenza specific CD8 T cells, the same stimulation experiment was performed using influenza derived NP and PA peptides. Interestingly, upon peptide stimulation, the CD49a subset displayed the highest frequencies of cytokine positive cells, consistent with differences in NP/PA specificity between the subsets (S Figure 7). CD49a cells showed significantly higher levels of both IFNγ and TNF compared with DN, albeit at a lower overall response compared with anti-CD3/CD28 stimulation (S Figure 7). Of note, IL-17A, IL-21, and IL-10 were also examined within these experiments, however, no protein staining was detected under the stimulation conditions described.

**Figure 5.**
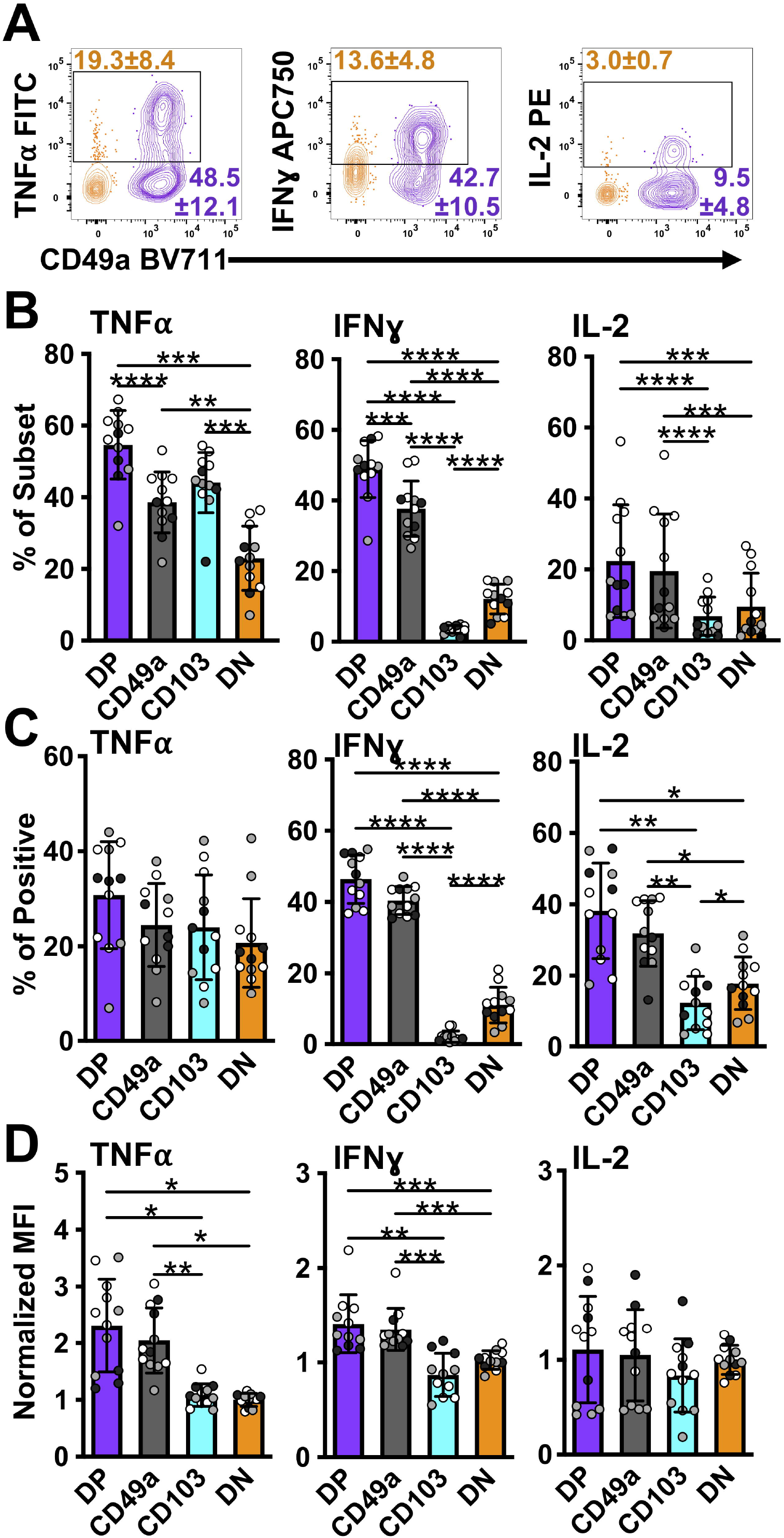
DP and CD49a CD8 memory T cells express a higher frequency of antiviral and survival proteins. Representative flow cytometry plots showing DP cells compared with DN cells after 6 hours of *in vitro* stimulation (A). Percentage of each subset positive for the cytokine (B) and percentage within the cytokine positive populations (C). MFI normalized to DN (D). Data for B-D shown as mean with standard deviation and each individual sample. *p<0.05 **p<0.01 ***p<0.001 ****p<0.0001 based on a repeated measures one-way ANOVA with Greenhouse-Geisser correction, followed by *post-hoc* testing comparing all groups through Tukey’s multiple comparisons test. Data is three experiments combined with black, grey, or white indicating the individual experiment. n=12 total DP CD49a CD103 DN

T_RM_ have previously been said to orchestrate both innate and adaptive responses, however, the mechanisms controlling these outcomes were not fully clear [6] [7]. Supported by the RNAseq data, we hypothesized that T_RM_ have the capacity to release both chemokines and growth factors to recruit and aid in APC maturation. To determine the breadth of the chemokine response of memory CD8 T cells, CCL3 and CXCL9 were analyzed in addition to CCL1. All of these chemokine genes were upregulated in DP compared with DN at the transcript level (Table 2). As suggested by clustering, CCL1 was expressed only in CD49a and DP cells, indicating a unique function of CD49a positive subsets, though these were more frequently found in DP cells than in CD49a (Figure 6A-C). This was consistent with results from NP and PA peptide stimulation (S Figure 7). CCL3 was expressed at the highest frequency, overall contribution, and MFI in DP cells and a higher percentage in CD49a versus CD103 and DN (Figure 6A-D). Additionally, the growth factor GM-CSF was expressed predominantly in CD49a expressing cells (DP and CD49a), showing both increased percentages within the subsets as well as increased contribution to the GM-CSF positive population (Figure 6A-C). We were unable to detect CXCL9 protein within the CD8 T cells using this re-stimulation approach. Overall, this data paints a picture of polyfunctional CD49a expressing cells as signaling hubs, both directing and contributing to an effective antiviral response.

**Figure 6.**
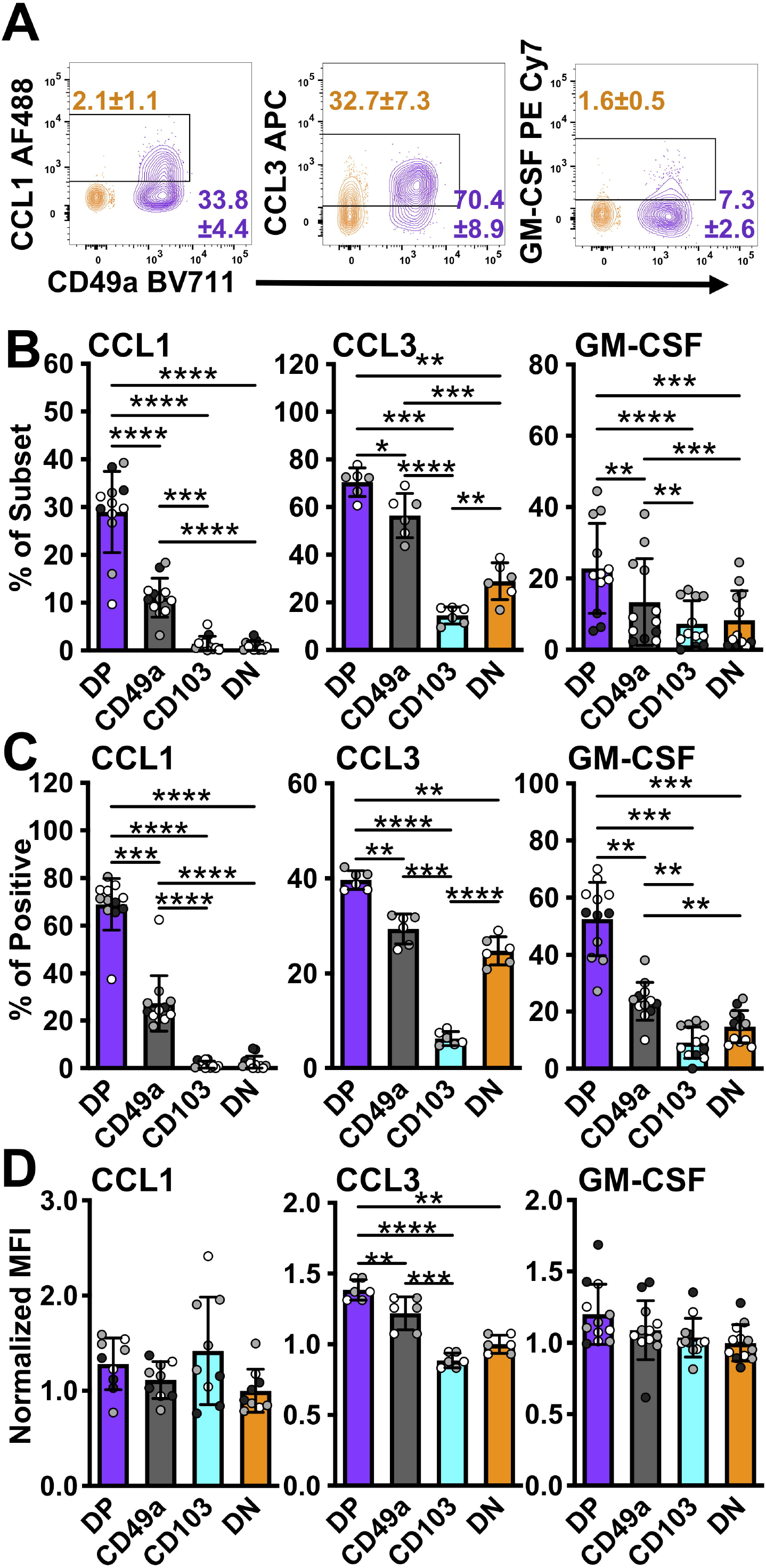
DP and CD49a CD8 memory T cells express higher frequencies of chemokine and growth factor producing cells. Representative flow cytometry plots showing DP cells compared with DN cells after 6 hours of stimulation *in vitro* stimulation (A). Percentage of each subset positive for the chemokine (B) and percentage within the chemokine positive populations (C). MFI normalized to DN (D Data for B-D shown as mean with standard deviation and each individual sample. *p<0.05 **p<0.01 ***p<0.001 ****p<0.0001 based on a repeated measures one-way ANOVA with Greenhouse-Geisser correction, followed by *post-hoc* testing comparing all groups through Tukey’s multiple comparisons test. Data is two (CCL3) or three experiments combined with black, grey, or white indicating the individual experiment. n=6 for CCL3 n=12 for CCL1 and GM-CSF DP CD49a CD103 DN

In addition to cytokines and chemokines, the increase in transcript levels of cytotoxic-associated molecules suggested that T_RM_ retain the ability to kill target cells (Table 2). DP cells display increases in mRNA for granzymes A, B, and C, and perforin. Furthermore, transcript for apoptosis-inducing FasL and TRAIL were increased (Table 2) [51; 52]. At 6 hours post-stimulation, memory CD8 T cells were evaluated for Lamp-1 staining, an indicator of vesicle:plasma membrane fusion, granzymes A, B, and C, perforin, FasL, and TRAIL [53]. Even at 6 hours post-stimulation, low levels of granzyme A and perforin were detected in DP cells with higher frequencies than in DN cells (Figure 7A). TRAIL and granzymes B and C were not detected at this time point (S Figure 8 and data not shown). While this was consistent with the idea that the function of T_RM_ is to release IFNγ rather than kill infected targets, it conflicted with mRNA levels [54] [49]. To further complicate our understanding, over half of the cells were surface Lamp-1 positive (Figure 7A). This suggested that despite the addition of transport inhibitors, vesicular fusion still occurred.

**Figure 7.**
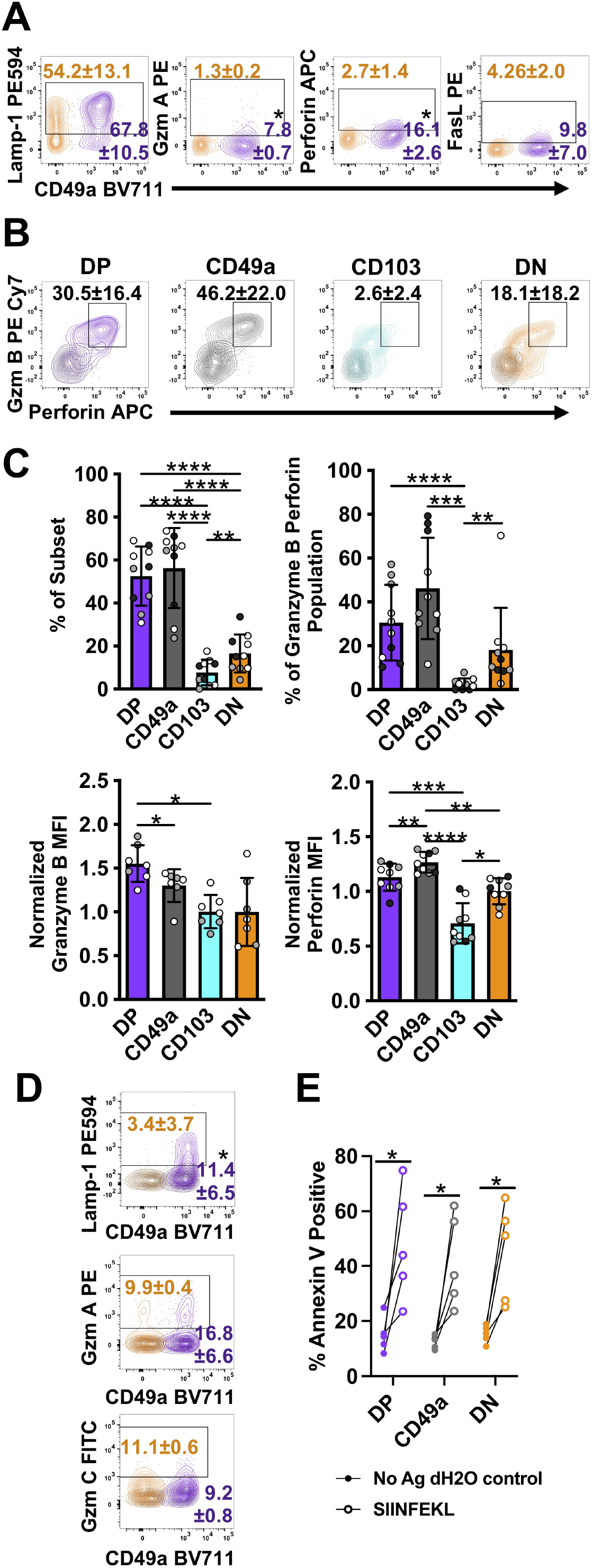
DP and CD49a CD8 memory T cells express a higher frequency cytolytic granules. Representative flow cytometry plots showing DP cells compared with DN cells after 6 hours of *in* vitro stimulation (A) and 30 hours of stimulation with media control overlaid (B) (Mean and standard deviation). Mean of each subset and individual samples positive for granzyme B and perforin (C) and percentage within the granzyme B/perforin double positive population (C). MFI normalized to DN for granzyme B and perforin (C). Comparison of DP and DN cells after 24 hours of stimulation with the mean and standard deviation of the percentage shown. Media control is overlaid in darker shade of the subset color (D). Percentage of Annexin V+ EL4 cells with vehicle control or antigen loading after 13 hour co-culture with T cell subsets sorted from lung tissue CD8 T cells at day 21+ post-infection. * p<0.05 based on a paired T test on one experiment representative of 3 independent experiments (A,D), p<0.05 based on paired T test on two combined experiments (E), or *p<0.05 **p<0.01 ***p<0.001 ****p<0.0001 based on a repeated measures one-way ANOVA with Greenhouse-Geisser correction, followed by *post-hoc* testing comparing all groups through Tukey’s multiple comparisons test (B,C) of three experiments combined with black, grey, or white indicating the individual experiment. DP CD49a CD103 DN

It is possible that T_RM_ cells may be pre-loaded with cytolytic granules that are not blocked from degranulation by ER and Golgi transport inhibitors. Furthermore, in agreement with a paper examining human skin T_RM_ cells, the inability to detect granzymes and high levels of perforin may be due to altered kinetics with increased time necessary for new granules to be formed (S Figure 8A)[13]. To test this, cells were incubated for either 12 (data not shown) or 24 hours prior to the addition of transport inhibitors. To stain for degranulation, αLamp-1 was added 2 hours prior to addition of transport inhibitors. Cells were then incubated for an additional 6 hours after addition of the inhibitors, and subsequently stained for granzymes and perforin. Using this protocol, CD49a expressing subsets from the lungs and the airways produced dramatic levels of perforin and granzymes (Figure 7B,C and S Figure 8B,C). In fact, CD49a expressed higher levels perforin per cell compared with DN cells, validating the RNAseq results. Not surprisingly, at this time point, DP cells expressed a higher proportion of Lamp-1 stained cells compared with DN, however, no increase in either granzymes A or C was observed (Figure 7D). Of note, while skin T_RM_ required only IL-15 in culture to produce granzyme B and this effect was additive in the presence of TCR stimulation, exogenous cytokines had no additional effect on the lung CD8 T cells evaluated in this model, and their addition in the absence of TCR stimulation was insufficient to induce cytotoxicity (data not shown) [13]. To determine whether granzyme B and perforin were indicative of actual killing ability, an *in vitro* killing assay was set up using EL4 cells as APCs. To minimize the known differences in tetramer positivity between the different subsets, the transgenic OT-I adoptive transfer model was utilized [55] [56]. Briefly, one day prior to infection with a SIINFEKL expressing HKx31 influenza A variant, GFP OT-I T cells were transferred intravenously. At greater than 21 days post-infection, lung tissue CD8 T cells were sorted based on their integrin phenotype. In this model, CD103 single positive cells were almost non-existent, so only DP, CD49a, and DN were evaluated. DP, CD49a, and DN cells were cultured with SIINFEKL pulsed Cell Tracker Violet labeled EL4 cells or unloaded Cell Tracker Violet labeled vehicle control EL4 cells. After 13 hours of co-culture, cells were harvested and stained for Annexin 5. All subsets examined displayed increased frequencies of Annexin V positive target cells in the presence of peptide, compared with control cells (Figure 7E). Despite the fact that a higher frequency of DP and CD49a populations produced perforin and granzyme B compared with DN, there was no significant difference in the contribution to the population, supporting that both resident and circulating CD8 T cells can kill infected targets (Figures 7C,E). Taken together, the data suggests that T_RM_ cells may respond in sequence, first limiting the spread of infection through release of antiviral cytokines and recruitment of innate cells, followed by direct elimination of residual virally infected cells. It also suggests that the heterogenous populations of T_RM_, may specialize in discrete responses.

## Discussion

Memory CD8 T cells arise in response to pathogen exposure and are maintained in different locations throughout the body [3] [4] [5]. The memory cells can be defined by their ability circulate or persist as resident memory at the site of infection. T_RM_ in different organs have discrete requirements for development, maintenance, and the level of effector capacity [57]. The requirement for varied responses in the periphery is likely driven in part by the type of invading pathogen and local inflammatory milieu. This study aimed to interrogate the memory population(s) at a single peripheral site, and ask whether this diversity was still maintained. CD8 T cells in the lung tissue were examined after resolution of influenza A virus infection, with the hypothesis that the cells referred to as T_RM_ actually present a heterogenous mix of populations which have particular effector capabilities. Surprisingly, the memory CD8 T cell composition in the lung tissue was comprised not only of T_RM_ and T_EM_, but also cells which matched the phenotype of T_CM_. While it cannot be discounted that the approach used captured cells within lymphatics of the lungs, the cells within the airway (through bronchoalveolar lavage), contained similar populations. This suggests that even on the surface, the diversity of the CD8 memory T cells found in lung tissue is much higher than previously appreciated. Within the cells of T_RM_ phenotype, the canonical CD103 CD49a CD69 population was identified, in addition to other subsets which express only one of the integrins or lack CD69. Unexpectedly, CX3CR1 expressing cells were found within both CD103/CD49a cells and the CD49a single positive populations. Similar to what is observed within skin draining DCs, we propose that CX3CR1 on these cells may explain the small subset of cells T_RM_ found within lymphatics and the draining lymph nodes [58] [37]. Since CX3CL1 can also be expressed by epithelial and endothelial cells, it is also conceivable that receptor expression improves homing to infected cells or reentry into the vasculature [38].

The phenotypic differences observed may be due to the microenvironmental niches in which the cells reside – changes in TGFβ levels alone could account for integrin expression alterations [29]. Other studies have shown that the requirement for CD69 is not present in every organ, however, it is conceivable that it may be more critical for cells in close proximity to afferent lymphatics, than for cells embedded within the epithelial cell layer [15; 22]. Similarly, a requirement for CD103:E-cadherin interactions would not exist for a T cell within the parenchyma or migrating along the basement membrane. These findings set up and support the idea that the surface phenotype may be driven by the local microenvironment, and may in turn promote different responses and cell fate within that niche.

It was unclear, however, whether the subsets were truly distinct, or if the cells examined just represent plasticity or phenotypic intermediates. At baseline, the cells phenotypically defined as T_RM_ (DP and CD49a) examined directly *ex vivo* have discrete transcriptional profiles compared with non-CD49a expressing population. In this homeostatic state, the T_RM_ cells contained higher levels of mRNA for effector genes, however, minimal protein, if any was observed in the absence of stimulation. Although less intensively studied in T_RM_ cells, memory CD8 T cells have been shown to employ different mechanisms through which they repress the translation of accumulated mRNA[46; 59; 60]. While the specific processes underlying these observations warrant further research, we hypothesized that the amassed effector gene transcripts provide the ability for T_RM_ cells to rapidly respond upon reactivation.

After *in vitro* restimulation, all four subsets were transcriptionally distinct, and this held true when PCA was performed excluding the genes for CD49a and CD103. Cells expressing both CD103 and CD49a had the highest levels of effector responses overall, spanning antiviral cytokines, T-cell survival genes, chemokines, a growth factor, and cytolytic mediators. CD49a single positive cells, which are intermediate in effector function between DP and DN cells, actually displayed increased enrichment for genes associated with cytotoxicity. Evaluation of protein levels confirmed the majority of the transcript data after *in vitro* reactivation, however, the kinetics for some of these responses were delayed. We hypothesized that these cells may be pre-programmed to respond in sequence, facilitating viral clearance and recruitment of cells with APC potential, without destroying the epithelial barrier.

Consistent with many other studies, within 6 hours of TCR stimulation, DP and CD49a cells produce copious amounts of protective IFNγ [49]. These same populations of cells also secrete T-cell survival factor, IL-2. These immediate responses may enhance the lifespan of the T_RM_ cells or promote proliferation after rechallenge and clear virus with less damage to the airway epithelium.

Within this same time frame, DP cells and CD49a cells release chemokines CCL1 and CCL3 and the growth factor GM-CSF. Interestingly, cells that traffic from the spleen after influenza infection have previously been shown to produce CCL1 and GM-CSF, so it is intriguing that this profile of protection is conserved within the cells already present within the lungs[61]. The receptor for CCL1 (CCR8) is expressed on specific subsets of T_RM_, so this mechanism could be in place to recruit T_RM_ at other regions of the tissue to specific niches of infection[62]. Concurrently, CCL1 can recruit innate immune cells (myeloid and DC lineages) to the site of infection [62; 63]. GM-CSF has roles in both maturation of myeloid and APC populations, but also in the repair of the airway epithelium [64; 65; 66]. This strengthens the hypothesis that T_RM_ aim to clear infection in the absence of extensive tissue damage.

In line with this, despite high transcript levels for cytolytic mediators granzyme B and perforin, minimal protein can be detected at 6 hours post-stimulation. These data are consistent with the concept that T_RM_ are initially noncytolytic and that control of infection is primarily through release of antiviral molecules. However, similar to T_RM_ in other systems, stimulating the cells for longer time periods results for the accumulation of high levels of granzyme B and perforin and killing of target cells [49] [13]. Interestingly, both CD49a single positive and DP cells have comparable positive frequencies for these cytotoxic molecules, but CD49a cells produce higher levels of perforin, and demonstrate a much higher frequency after stimulation with NP/PA peptide. This suggests that during reinfection with a heterosubtypic virus, the CD49a cells may be more predisposed to killing target cells than their DP counterparts.

Overall, this study identified a previously unappreciated level of heterogeneity within the lung CD8 memory subset, even within the populations of cells labelled as T_RM_. Cells expressing CD49a display the highest levels of effector function, and the integrin phenotype may further identify cells capable of cytotoxicity and repair mechanisms. In another study CD49a expression was utilized to sort for downstream RNA sequencing, all integrin positive cells were evaluated, rather than only antigen experienced CD8 T cells. This difference alone makes it difficult to directly compare with the results from this study. However, despite that caveat, they found comparable results in regard to transcripts related to cytotoxicity (*GZMB* and *PRF1*), chemokines (*CCL4* and *CCL5*), and antiviral cytokines including *IFNG* [13].

With the knowledge that CD49a similarly defines polyfunctional T_RM_ in human skin and tumor infiltrating lymphocytes in melanoma, and supports cell motility, we propose that the ubiquity of collagen IV in the basement membrane of epithelial surfaces and the data presented here, further substantiate a probable role for CD49a in many, if not all T_RM_ populations [13] [27]. As a whole, the memory T cells in the lungs have the capacity to generate antiviral immunity, support T cell survival, enhance recruitment and maturation of other cells to the site of infection, with the potential to aid in repair of the tissue after clearance of the pathogen.

## Data availability statement

The data that support the findings of this study are available from the corresponding authors upon reasonable request. The RNAseq data is available through GEO Accession number GSE179653.

## Code availability statement

All code used for data analysis in this manuscript is available at https://github.com/tophamlab20.

## Acknowledgments

This work was made possible with the support and contributions of the University of Rochester Medical Center Genomics Research Center (GRC), Flow Cytometry Core, and the Vivarium as well as the NIH Tetramer Core at Emory University.

## Notes

https://www.ncbi.nlm.nih.gov/geo/query/acc.cgi?acc=GSE179653

### Competing Interest Statement

The authors have declared no competing interest.

