## Supplemental Figures S1-S8 for "CD49a Identifies Polyfunctional Memory CD8 T cell Subsets that Persist in the Lungs after Influenza Infection"

### S Figure 1

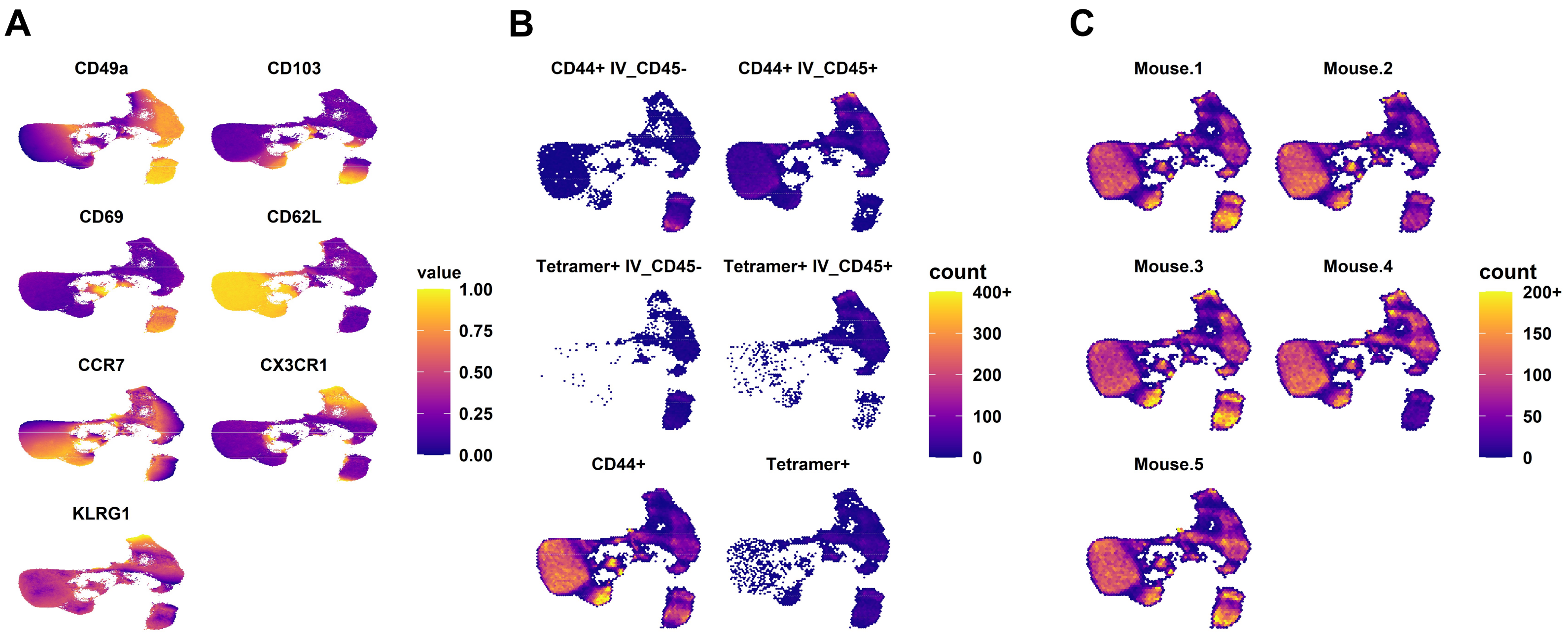


**S Figure 1. UMAPs to show surface marker expression across tissues, activated and influenza specific populations, and consistency across mice.** Surface marker expression from concatenated blood, spleen, MLN, BAL, lung vasculature, and lung tissue samples (A). CD8 T cells separated further by IV labeling, CD44, and tetramer staining (B). Individual mice (C).

#### S Figure 2

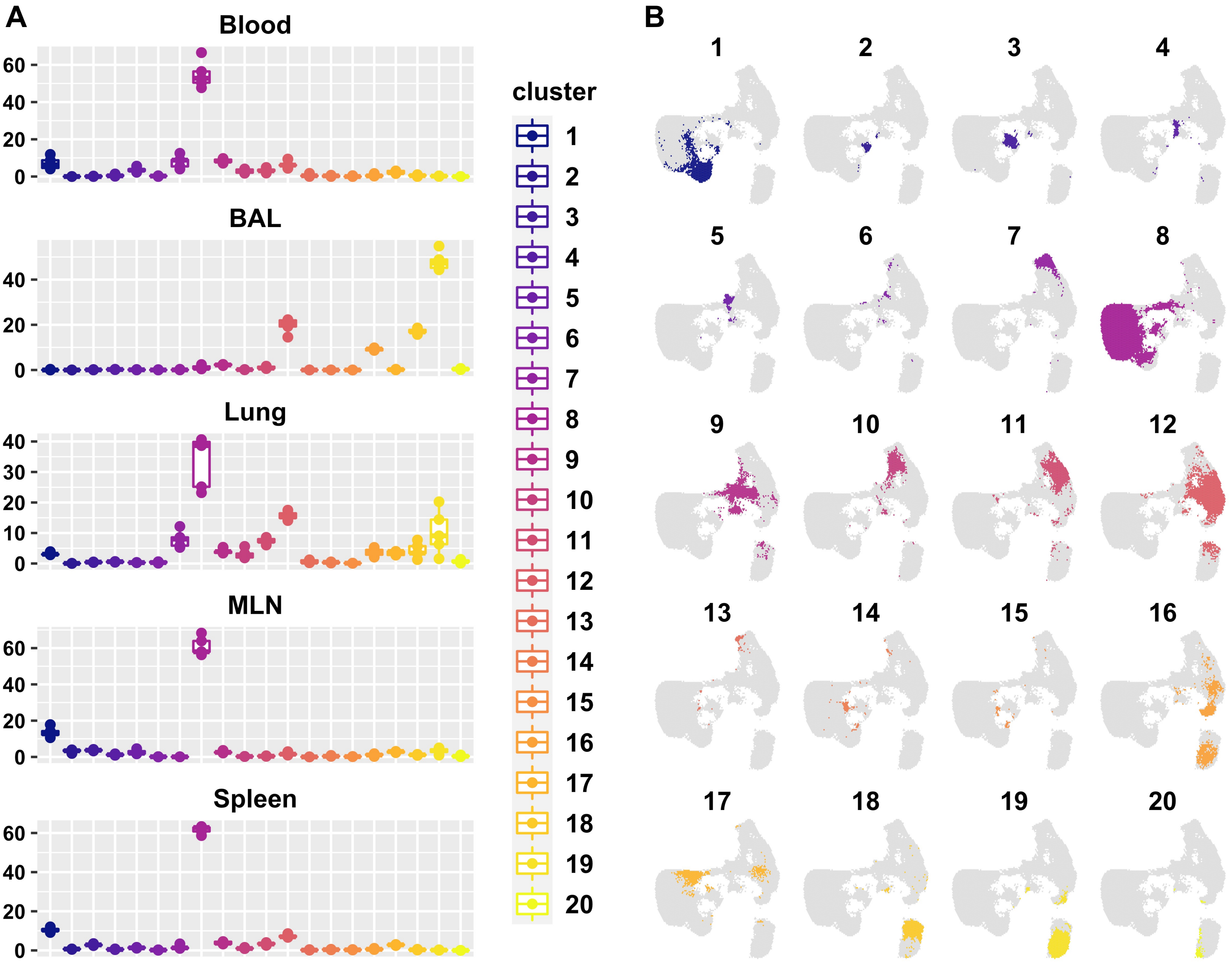


**S Figure 2. Cluster frequency across organs and UMAPs faceted by cluster.** Box plot frequencies shown for each cluster in separate organs (A). Individual clusters projected on to the UMAP (B).

### S Figure 3

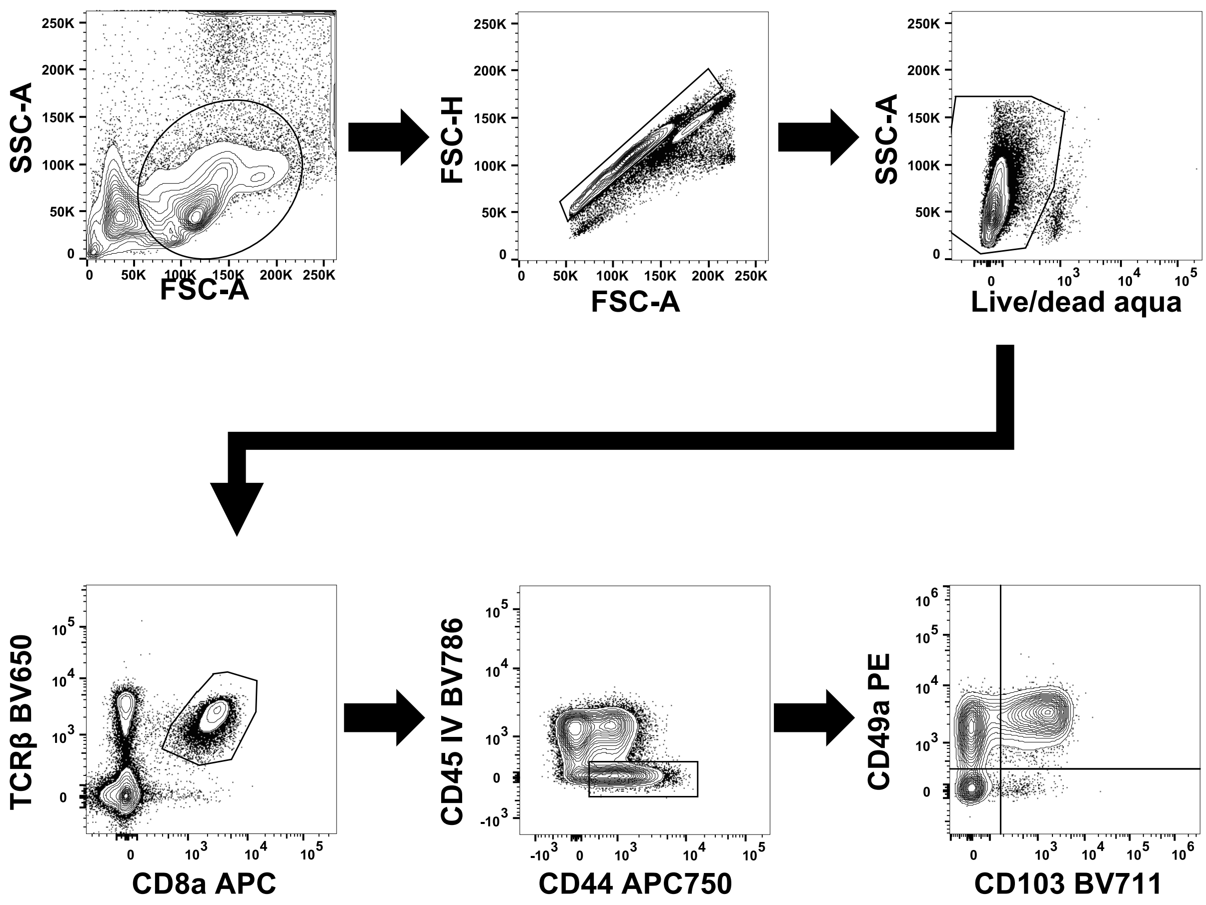


**S Figure 3. Gating strategy.** A liberal lymphocyte gate based on SSC-A and FSC-A was applied, followed by doublet exclusion by FSC height and area parameters. Cells that stained for live/dead aqua stain were excluded, and CD8 T cells were identified by expression of TCRβ and CD8a. CD8 T cells were further selected for CD44 and negative for intravenous (IV) labeling with CD45. These cells were sorted based on CD49a and CD103 profile into DP, CD49a, CD103, and DN populations. Of note, for previously *in vitro* activated cells, TCRβ was downregulated and all CD8+ cells were included.

**S Figure 4**

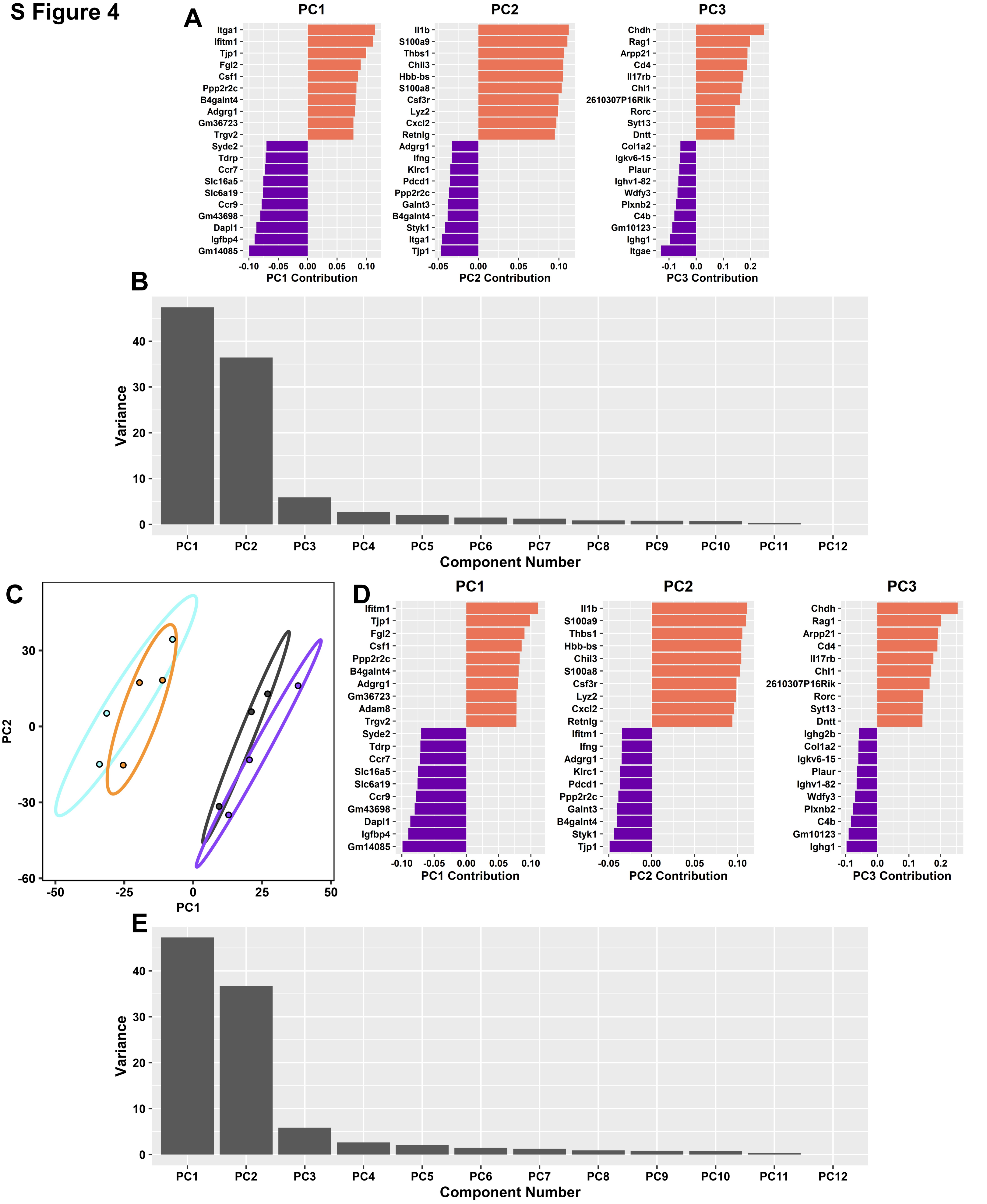


**S Figure 4. Contribution to PCA.** The top loading genes driving the discrimination of PC1, PC2, and PC3 in cells directly *ex vivo* are shown are shown (A) as well as the component contribution to variance (B). PCA performed without CD49a and CD103 (C) and the top loading genes (D) and variance (E).

#### S Figure 5

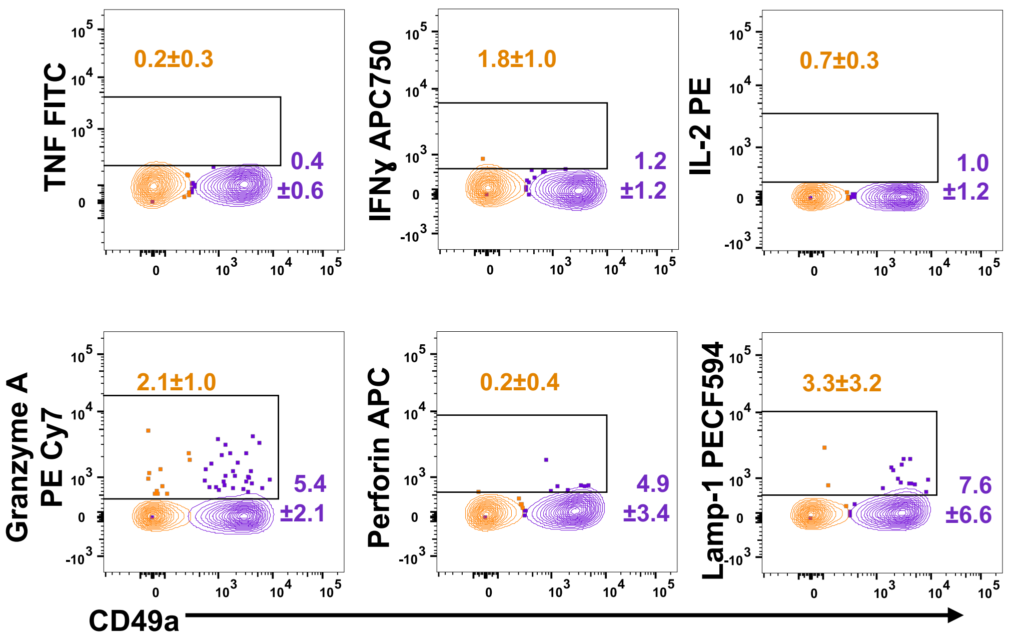


**S Figure 5. DP cells do not produce TNF, IFNɣ, or IL-2 in the absence of stimulation, but do express low percentages of granzyme A, perforin, and Lamp-1.** Representative flow cytometry plots showing the percentage and standard deviation of positive cells for DP and DN populations.

**S Figure 6**

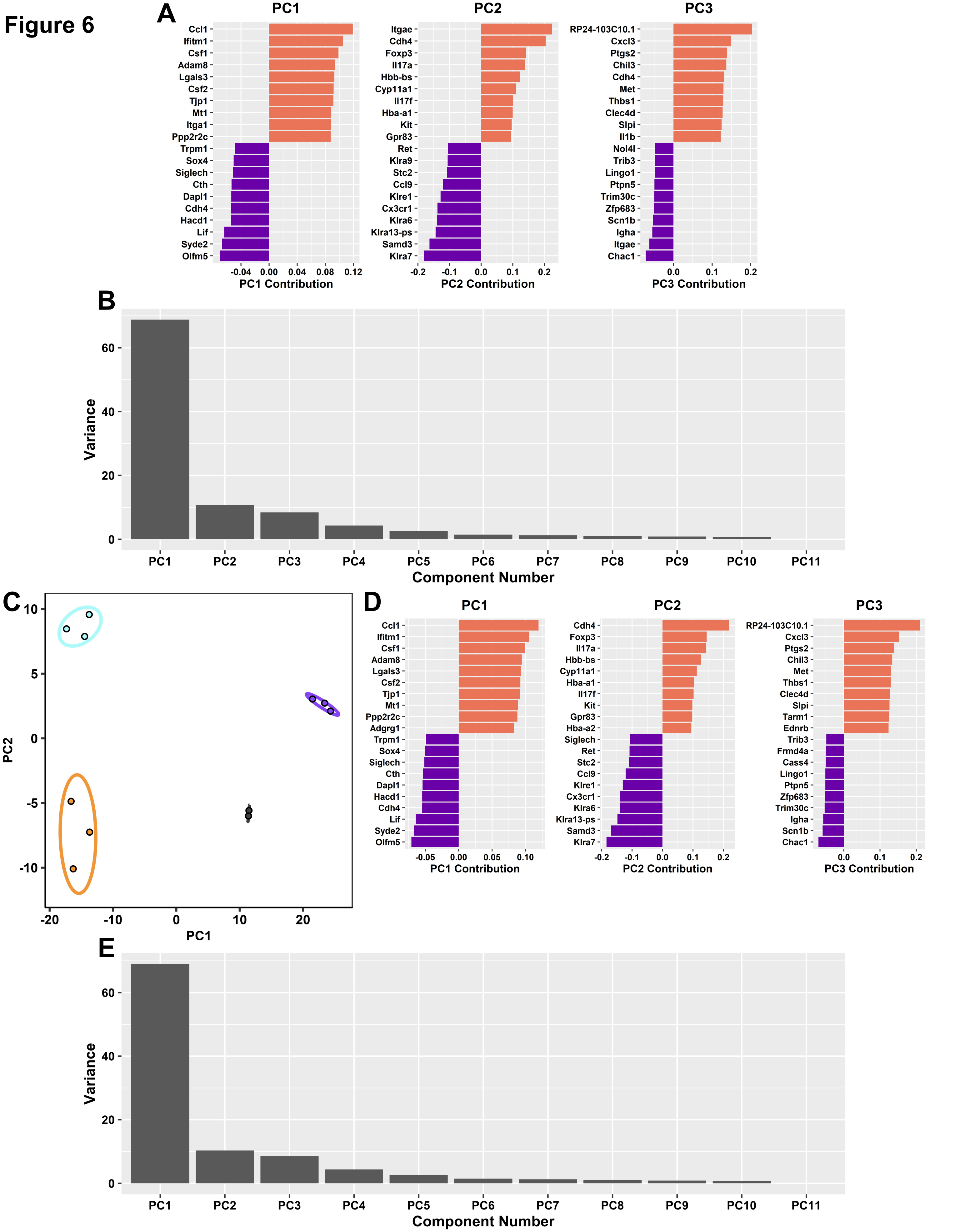


**S Figure 6. Contribution to PCA.** The top loading genes driving the discrimination of PC1, PC2, and PC3 in cells after *in vitro* stimulation are shown (A) as well as the component contribution to variance (B). PCA performed without CD49a and CD103 (C) and the top loading genes (D) and variance (E).

### S Figure 7

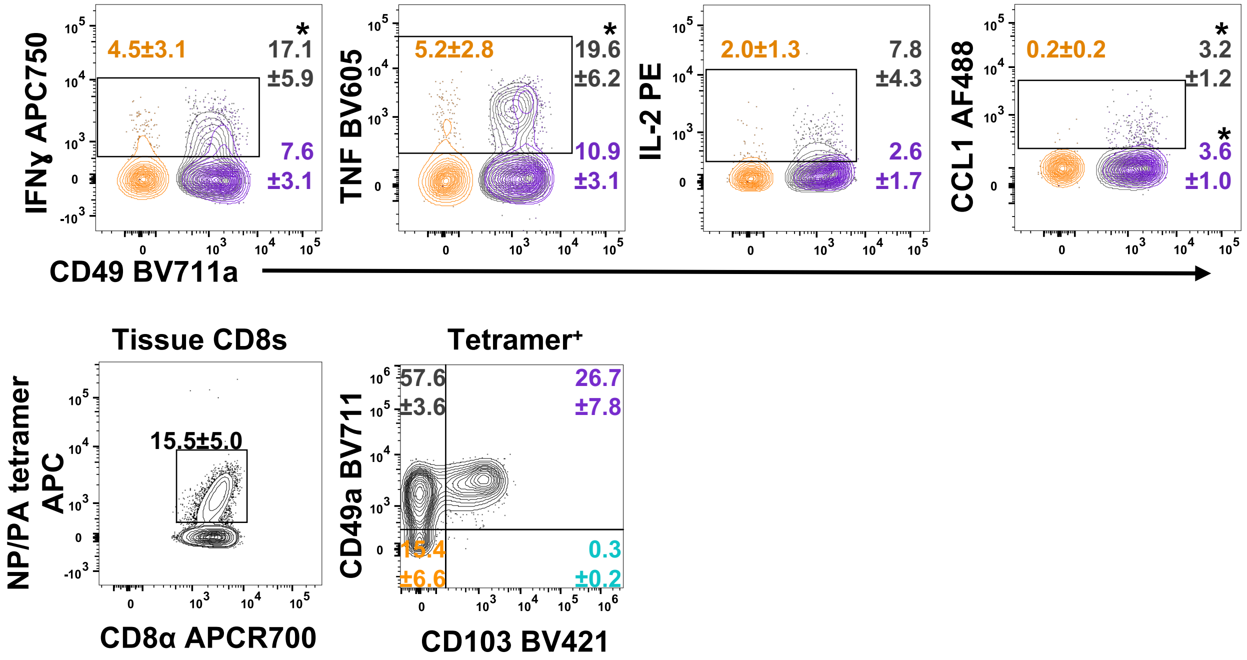


**S Figure 7. CD49a cells show increased IFNɣ, TNF, and CCL1 responses after NP and PA peptide stimulation.** Representative flow cytometry plots showing the DP, CD49a, and DN populations after a 6 hour stimulation with NP and PA peptides derived from HKx31 influenza virus and the positive frequency and standard deviation for IFNɣ, TNF, IL-2, and CCL1. * $p<0.05$ based on a repeated measures one-way ANOVA with Greenhouse-Geisser correction, followed by *post-hoc* testing comparing groups to DN using Dunnett's multiple comparisons test. Data are from one experiment representative of two independent experiments with an n≥3 mice/experiment.

S Figure 8

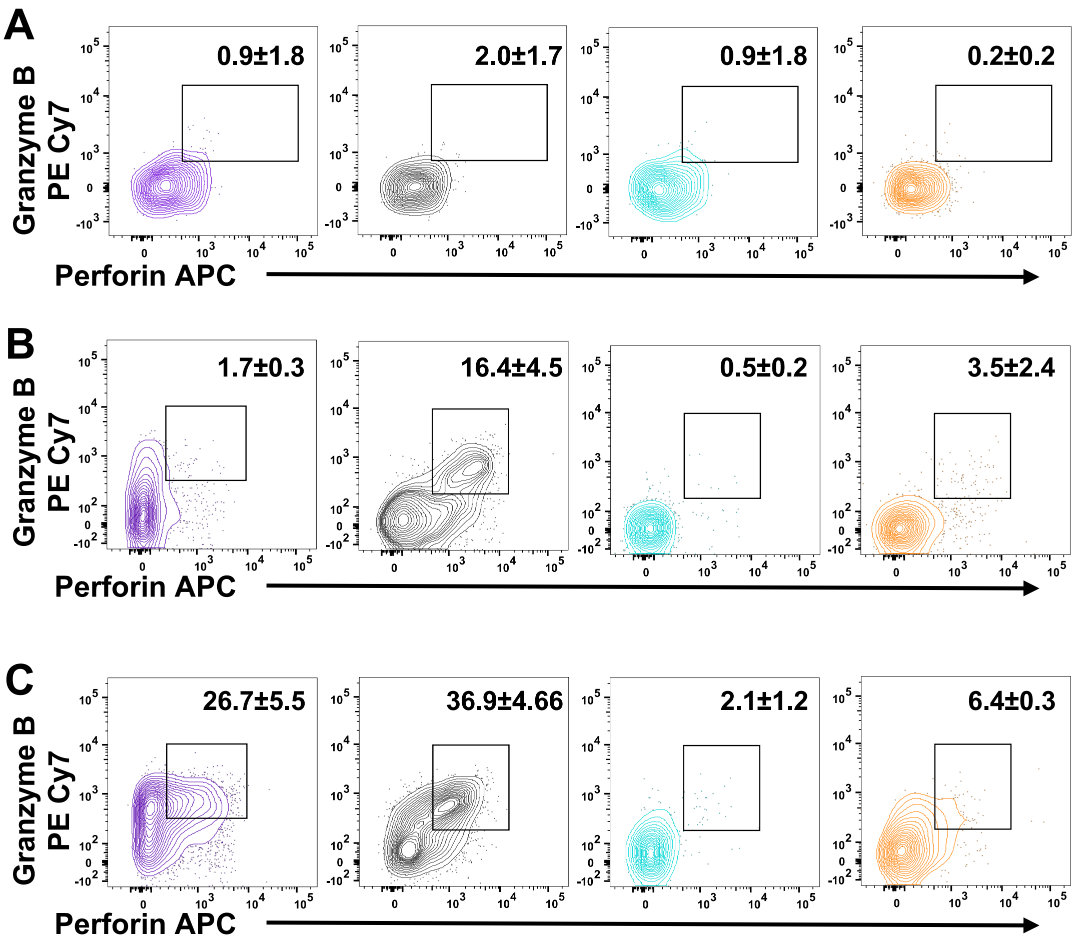


**S Figure 8. Perforin and granzyme B production.** Granzyme B and perforin staining of lung CD8 T cells after 6hrs of stimulation with αCD3/28 (A). Lung CD8 T cells after 30hrs of stimulation with NP/PA peptide (B). CD8 T cells from the airways were stimulated with αCD3/28 for 30hrs and stained for granzyme B and perforin (C). **DP** **CD49a** **CD103** **DN**
