## Supplemental Table 1 for "CD49a Identifies Polyfunctional Memory CD8 T cell Subsets that Persist in the Lungs after Influenza Infection"

**S Table 1. Antibodies.** Antibodies that were used within the experiments in this paper are indicated with the company, the clone, and the fluorochrome(s).

| **Antigen** | **Company** | **Clone** | **Fluorochrome** |
| --- | --- | --- | --- |
| CCL1 | Invitrogen | 148113 | AF488 Click-IT^TM^ |
| CCL3 | Invitrogen | 39624 | APC |
| CD49a | BD Biosciences | Ha31/8 | Brilliant Violet^TM^ 711 or PE |
| IL-2 | BD Biosciences | JES6-5H4 | PE |
| CD8a | BD Biosciences | 53-6.7 | APC-R700 |
| CD103 | Biolegend | 2E7 | Brilliant Violet^TM^ 421, 711, or PE |
| TCRB | Biolegend | H57/597 | Brilliant Violet^TM^ 650 |
| CD45 | Biolegend | 30-F11 | Brilliant Violet^TM^ 785 |
| TNFa | Biolegend | MP6-XT22 | Brilliant Violet^TM^ 605 or FITC |
| CD69 | Biolegend | H1.2F3 | PE/Dazzle^TM^ 594 |
| FasL | Biolegend | MFL3 | PE |
| TRAIL | Biolegend | N2B2 | PE Cy7 |
| Granyme A | Biolegend | 3G8.5 | PE or PE Cy7 |
| Granzyme B | Biolegend | QA16A02 | PE Cy7 |
| Granzyme C | Biolegend | SFC1D8 | FITC |
| Perforin | Biolegend | S16009A | APC |
| Lamp-1 | Biolegend | 1D4B | PE/Dazzle^TM^ 594 |
| CD44 | Biolegend | IM7 | APC/Fire^TM^ 750 |
| IFNg | Biolegend | XMG1.2 | APC/Fire^TM^ 750 |
| GM-CSF | Biolegend | MP1-22E9 | PE Cy7 |
| CD62L | Biolegend | MEL-14 | PE Cy7 |
| CX3CR1 | Biolegend | SA011F11 | Brilliant Violet^TM^ 605 |
| KLRG1 | Biolegend | 2F1/KLRG1 | FITC |
| CCR7 | Biolegend | 2B12 | PE |
| CD8a | Biolegend | 53-6.7 | APC |
