## Supplemental Table 2 for "CD49a Identifies Polyfunctional Memory CD8 T cell Subsets that Persist in the Lungs after Influenza Infection"

**S Table 2. Cluster statistics for Figure 1.**

| **Cluster** | **DP vs. CD49a** | **DP vs. CD103** | **DP vs. DN** | **CD49a vs. CD103** | **CD49a vs. DN** | **CD103 vs. DN** |
| --- | --- | --- | --- | --- | --- | --- |
| **1 ****** | ns | **** | ns | **** | n/a | **** |
| **2 *** | ns | ns | ns | ns | ns | ns |
| **3 ****** | n/a | ns | ** | ns | ** | ** |
| **4 ****** | n/a | **** | n/a | **** | n/a | **** |
| **5 **** | n/a | ns | ** | ns | ** | ns |
| **6 **** | n/a | ns | ** | ns | ** | ns |
| **7 ****** | ** | ns | ** | *** | ns | *** |
| **8 ****** | n/a | ** | **** | ** | **** | *** |
| **9 ***** | ns | ** | **** | * | **** | * |
| **10 **** | n/a | ns | **** | ns | **** | * |
| **11 ****** | **** | ns | *** | *** | ** | ** |
| **12 ****** | **** | * | ** | *** | ** | * |
| **13 *** | ns | n/a | ns | ns | ns | ns |
| **14 ^ns^** | n/a | n/a | n/a | n/a | n/a | n/a |
| **15 ^ns^** | n/a | n/a | n/a | n/a | n/a | n/a |
| **16 ****** | *** | ** | **** | ** | ** | *** |
| **17 ***** | ** | ns | **** | ** | ns | **** |
| **18 ****** | **** | ns | ns | *** | **** | ns |
| **19 ****** | **** | * | **** | *** | ** | *** |
| **20 ***** | **** | * | **** | ns | n/a | ns |
