## Supplemental Table 7 for "CD49a Identifies Polyfunctional Memory CD8 T cell Subsets that Persist in the Lungs after Influenza Infection"

**S Table 7. Cluster statistics for Figure 4.**

| **Cluster** | **DP vs. CD49a** | **DP vs. CD103** | **DP vs. DN** | **CD49a vs. CD103** | **CD49a vs. DN** | **CD103 vs. DN** |
| --- | --- | --- | --- | --- | --- | --- |
| **1****** | * | ** | ** | ** | ** | ns |
| **2*** | ns | * | * | ns | ns | ns |
| **3**** | ns | ** | ns | * | ns | * |
| **4**** | * | ** | ns | *** | ns | * |
| **5*** | * | ns | ns | ns | ns | * |
| **6ns** | n/a | n/a | n/a | n/a | n/a | n/a |
| **7**** | * | * | ** | * | * | ns |
| **8****** | ns | ** | ** | ** | ** | * |
| **9***** | ns | ** | ** | ** | * | ns |
| **10*** | * | ns | * | ns | ns | ns |
| **11*** | ns | * | ns | * | ns | ns |
| **12****** | ns | *** | * | ** | * | ns |
| **13**** | * | ** | ** | ns | ns | ns |
| **14**** | ** | ** | ** | ns | ns | ns |
| **15ns** | n/a | n/a | n/a | n/a | n/a | n/a |
| **16**** | ns | ** | ** | ** | ** | ns |
| **17**** | ** | ** | ** | ns | * | ns |
| **18***** | * | ** | ** | * | * | n/a |
